# A *cis*-regulatory atlas in maize at single-cell resolution

**DOI:** 10.1101/2020.09.27.315499

**Authors:** Alexandre P. Marand, Zongliang Chen, Andrea Gallavotti, Robert J. Schmitz

## Abstract

*Cis*-regulatory elements (CREs) encode the genomic blueprints of spatiotemporal gene expression programs enabling highly specialized cell functions. To identify CREs at cell-type resolution in *Zea mays*, we implemented single-cell sequencing of Assay for Transposase Accessible Chromatin (scATAC-seq) in seedlings, embryonic roots, crown roots, axillary buds, and pistillate and staminate inflorescence. We describe 92 states of chromatin accessibility across 165,913 putative CREs and 52 known cell types. Patterns of transcription factor (TF) motif accessibility predicted cell identity with high accuracy, uncovered putative non-cell autonomous TFs, and revealed TF motifs underlying higher-order chromatin interactions. Comparison of maize and *Arabidopsis thaliana* developmental trajectories identified TF motifs with conserved patterns of accessibility. Cell type-specific CREs were enriched with enhancer activity, phenotype-associated genetic variants, and signatures of breeding-era selection. These data, along with companion software, *Socrates*, afford a comprehensive framework for understanding cellular heterogeneity, evolution, and *cis*-regulatory grammar of cell-type specification in a major crop.

## INTRODUCTION

Global consumption of maize per kilogram per person is expected to increase by 163% by 2050 (CIMMYT 2016). However, climate instability and disease strain are increasingly predicted to lower global maize yields by more than 10% over the same time frame (Tigchelaar et al., 2018; Zhao et al., 2017). As the foundational unit of plants, individual cells are responsible for the synthesis, transportation, and storage of rich primary and secondary metabolites that sequesters carbon and provide human nourishment. However, our understanding of cell-type functions in plants has been precluded by technical limitations imposed by the cell wall and an inability to culture homogenous cell lines, in contrast to mammalian models.

Past studies querying plant cell-type functions employed technically challenging experimental procedures to circumvent these obstacles, such as fluorescence-activated cell sorting (FACS) of GFP-tagged marker proteins, or isolation of nuclei tagged in specific cell types (INTACT) (Birnbaum et al., 2003; Brady et al., 2007; Deal and Henikoff, 2011). Although instrumental to understanding certain cell types, a shortcoming of these methods is the requirement of transgenesis and prior information regarding cell-type specificity for purification, thereby occluding unbiased efforts for discovery of unknown and poorly studied cell types. As a result, molecular profiling on a genome-wide scale of individual cells and cell types in plants have been largely limited to the roots of *Arabidopsis thaliana* and a handful of isolated tissues (Dorrity et al., 2020; Farmer et al., 2020; Jean-Baptiste et al., 2019; Lee et al., 2019; Lopez-Anido et al., 2020; Nelms and Walbot, 2019; Ryu et al., 2019; Shulse et al., 2019). Although critical for driving innovation in biotechnology, a comprehensive organismal cell-type atlas has yet to be realized in any plant species.

Development, differentiation, and response to environment in eukaryotic cells rely on precise spatiotemporal gene expression mediated by *cis*-regulatory elements (CREs) (Andersson and Sandelin, 2020; Cusanovich et al., 2018; Long et al., 2016; Lu et al., 2019; Marand et al., 2017; Teale et al., 2006; Wittkopp and Kalay, 2011). CREs encode DNA binding sites for transcription factors (TF) that cooperatively dictate transcriptional outcomes (Buchler et al., 2003; Cheng et al., 2012; Consortium, 2012; Gerstein et al., 2012; Ravasi et al., 2010). In metazoan genomes, CCCTC-binding factor (CTCF) directs higher-order chromatin interactions that facilitate spatial proximity of CREs and their target genes (Phillips and Corces, 2009). Plant genomes are also generalized by higher-order chromatin architecture, yet all plant lineages lack an ortholog to CTCF (Heger et al., 2012). How cells interpret the *cis-*regulatory code, establish diverse chromatin contact landscapes, and adopt specialized functions in discrete cell types are essential questions for understanding the rules governing biology. As a consequence of their centrality in establishing cell identity and function, a growing body of evidence point to genetic variation of CREs as a major source of phenotypic innovation, including disease and evolutionary divergence (Rebeiz and Tsiantis, 2017; Villar et al., 2015). However, it has become increasingly apparent that genetic variants may only affect CRE activity in a subset of cell types (Hekselman and Yeger-Lotem, 2020). We reasoned that a thorough investigation comprising the full spectrum of evolutionary changes through both inter- and intraspecies comparisons would be informative for detangling the cell type-specific contributions towards phenotypic variation.

Here, we describe the construction of a *cis*-regulatory atlas in the historically rich genetic model and crop species, *Zea mays*. We measure chromatin accessibility and nuclear gene expression in 72,090 cells across six major maize organs. Model-based normalization of chromatin accessibility enabled the identification and validation of diverse cell types, many of which lacked previous genome-wide characterization. We define the *cis*-regulatory combinatorial grammar underlying cell identity, reveal distinct TFs coordinating higher-order chromatin interactions, and demonstrate enhancer CREs with increased capacity for interactions as major contributors to phenotypic variation. Through an evolutionary lens, we uncover CREs and cell types targeted by modern breeding and evaluate the evolutionary impacts on *cis*-regulatory specification of cellular development between two highly diverged angiosperms (maize and *A. thaliana*). Finally, we present the R package “*Socrates*”, a unified framework for scATAC-seq pre-processing, normalization, downstream analysis, and integration with scRNA-seq data as a streamlined method for single-cell genomic studies.

## RESULTS

### Assembly and validation of a high-quality *ci*s-regulatory atlas in maize

To comprehensively assess *cis*-regulatory variation among cell types in a major crop species, we isolated nuclei using fluorescence-activated nuclei sorting (FANS) and generated single-cell chromatin accessibility profiles using Assay for Transposase Accessible Chromatin (scATAC-seq). Libraries were prepared from six major *Z. mays* L. cultivar B73 organs, including axillary buds, staminate and pistillate inflorescence, whole seedling (composed of stem, leaf, and coleoptile tissues), embryonic root tips, and post-embryonic crown roots as a representative sample of *cis-*regulatory diversity across a suite of maize cell types and tissues (**Figure 1A**, **S1A**, and **S1B**; **Table S1**).

**Figure 1.**
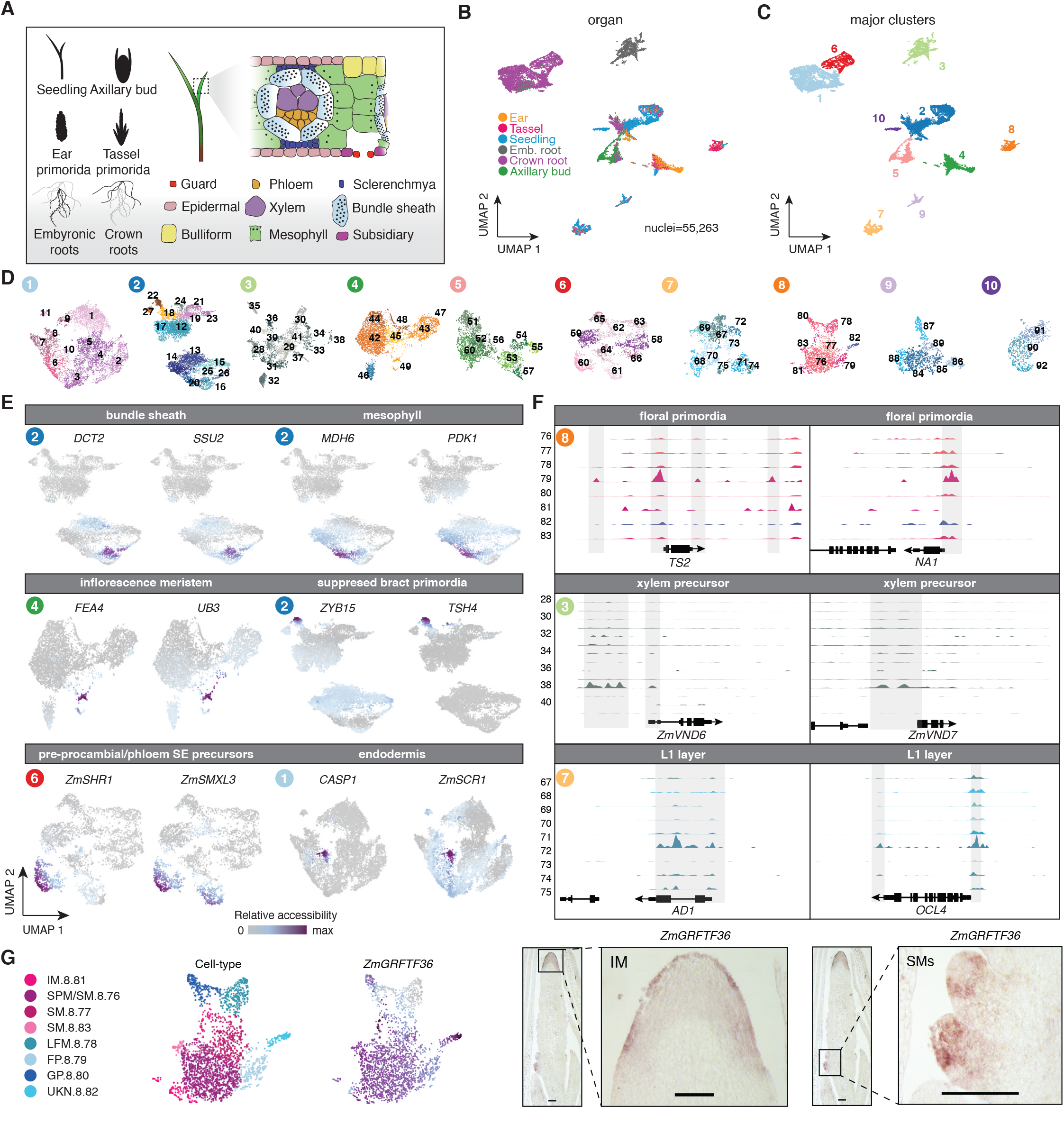
Atlas-scale cell type profiling from single nuclei chromatin accessibility in *Zea mays*. (A) Overview of experimental samples, with an example of the cell type diversity present in seedlings. (B) Nuclei similarity clustering as a UMAP embedding derived from the denoised quasibinomial Pearson’s residuals across all ACRs for each nucleus. UMAP embedding of nuclei colored by organ identity. (C) UMAP embedding of nuclei colored major cluster identity. (D) UMAP embedding of sub-cluster assignments following a second round of clustering within each major cluster. Sub-cluster color reflects the organ with the greatest proportion of nuclei in the cluster. See panel (B) for color code. (E) Cell type-specific enrichment of gene accessibility for a subset of marker genes associated with six different cell types. (F) Sub-cluster-specific chromatin accessibility profiles surrounding known marker genes for floral primordia, xylem precursors, and L1 epidermal cells. Bold, circled numbers indicate the cognate major cluster shown in panel c. Sub-cluster numeric identifications are present on the sides of the coverage plots. (G) Top, gene accessibility for *ZmGRFTF36*, an inflorescence and spikelet meristem enriched transcription factor with no previously known cell type-specificity. Bottom, RNA *in situ* hybridization of *ZmGRFTF36* in maize B73 staminate (tassel) primordia. FP, floral primordia; GP, glume primordia; IM, inflorescence meristem; LFM, lower floral meristem; SM, spikelet meristem; SPM, spikelet pair meristem; UKN, unknown.

We evaluated several quantitative and qualitative metrics reflective of high quality scATAC-seq data. In aggregate, scATAC-seq libraries were highly correlated between biological replicates and with previously generated organ-matched bulk ATAC-seq data (**Figure S1B**, **S2A**). Individual nuclei exhibited strong enrichment at transcription start sites (TSSs) (**Figure S1D**, **S2B** and **S2C**) and were consistent with the expected distributions of nucleosome-free and nucleosome-protected fragments (**Figure S1E** and **S2D**). By genotyping individual nuclei from a pooled population composed of B73 and Mo17 genotypes, we found 96% (4,944/5,177) were representative of a single genotype, validating our experimental approach (**Figure S1F–S1I**; **STAR Methods**). We identified a total of 56,575 nuclei passing quality filters (range: 4,704 - 18,393 nuclei per organ) with an average of 31,660 unique Tn5 integration sites per nucleus (**Figure S2C**, **S2E** and **S2F**; **Table S2**; **STAR Methods**).

Towards identifying clusters of nuclei resembling cell types, we first identified accessible chromatin regions (ACRs) by *in silico* sorting, resulting in a catalog of 165,913 putative CREs covering ∼4% of the maize genome (**Figure S3**; **STAR Methods**). To enable species-agnostic model-based analysis of scATAC-seq data, we developed an R package, termed ‘*Socrates’,* that streamlines data processing, clustering and downstream analysis. At its heart, ‘*Socrates*’ implements a regularized quasibinomial logistic regression framework to remove unwanted variation stemming from differences in nuclei read depth or other experimental factors (**Figure S4A**; **STAR Methods**). Following normalization with *Socrates*, we projected nuclei into a two-dimensional space using Uniform Manifold Approximation Projection (UMAP), revealing 10 major clusters unbiased by technical variation (**Figure 1B**, **1C**, and **S4B–S4E**).

Consistent with functional diversification of spatially distinct cells, most major clusters were generally composed of nuclei from the same organ (**Figure 1B** and **1C**). However, we also found evidence of common cell identities from nuclei in different organs, such as the co-localization of pistillate inflorescence, staminate inflorescence, and seedling nuclei in clusters 2, 4, 7 and 8 (**Figure 1B** and **1C**). Apparent heterogeneity within major clusters prompted us to implement a second round of partitioning for each major grouping, producing a total of 92 sub-clusters (hereafter referred to as clusters) with an average of 551 nuclei (**Figure 1D**; **STAR Methods**). Clear reproducibility and mitigation of technical variation in the UMAP embedding justifies ‘*Socrates’* as a robust approach for establishing shared cell identities across heterogenous organs through the removal of technical variation typical of single-cell experiments.

### Cell-type annotation and validation by *in situ* hybridization

To annotate clusters with corresponding cell types, we integrated chromatin accessibility information on a per-gene and nucleus basis as a proxy for gene expression (bulk RNA-seq versus aggregate scATAC-seq Spearman’s correlation coefficient = 0.54-0.58; **Figure S4F**; **STAR Methods**). We then (*i*) evaluated differential accessibility among clusters for a manually curated list of 221 literature-derived known marker genes (**Table S3**), (*ii*) classified cell types of individual nuclei with a multinomial logistic classifier trained on nuclei with discriminative cell type-specific signatures, and visually assessed (*iii*) accessibility scores of *a priori* marker genes over the UMAP embeddings and (*iv*) cluster-aggregated ATAC-seq coverages (**Figure 1E**, **1F**, **S5A**; **Table S4**; **STAR Methods**). Patterns of gene accessibility were consistent with *a priori* information of cell type/domain-specific expression, such as co-localized accessibility of bundle sheath-specific genes *DICARBOXYLIC ACID TRANSPORTER1* (*DCT2*) and *RIBULOSE BISPHOSPHATE CARBOXYLASE SMALL SUBUNIT2* (*SSU2*), and mesophyll-specific genes *MALATE DEHYDROGENASE6* (*MDH6*) and *PYRUVATE DEHYDOGENASE KINASE1* (*PDK1*), in addition to many other previously established cell type-specific marker genes (**Figure 1E**, **1F**, and **S5A**) (Chang et al., 2012).

To corroborate predicted cell-type annotations, we performed RNA *in situ* hybridization for a subset of differentially accessible genes with no prior evidence of cell type-specificity. In all cases (5/5), *in situ* expression patterns were in line with the predicted localization based on gene accessibility (**Figure 1G** and **S5B**). Estimates of cell-type proportions within and across organs were also concordant with prior observations, such as the occurrence of vascular bundle sheath and parenchymal mesophyll cells within multiple organs, including those derived from stem and leaf tissues of seedlings and within pistillate and staminate inflorescence (**Figure S5C**; **Table S4**) (Langdale et al., 1989).

### Integration of chromatin accessibility and gene expression from individual nuclei

To evaluate the correspondence between nuclear transcription and chromatin accessibility on a global scale, we sequenced the transcriptomes of 15,515 nuclei derived from 7-day old seedlings using single-nucleus RNA-seq (snRNA-seq; **STAR Methods**). We then integrated the corresponding nuclei with seedling-derived nuclei from scATAC-seq via integrative non-negative matrix factorization (iNMF; **Figure 2A**; **STAR Methods**) (Welch et al., 2019). Co-embedding nuclei on the basis of chromatin (n=11,882) and nuclear gene expression (n=15,515) revealed 19 clusters of nuclei with similar genome-wide profiles (**Figure 2B** and **2C**; **STAR Methods**). Comparison of the two modalities (n=36,322 genes) across clusters revealed a striking correspondence between the patterns of chromatin accessibility and nuclear transcription (Spearman’s correlation coefficient range across cell types = [0.52-0.69]; **Figure 2D**). The molecular relationship between chromatin accessibility and gene expression was further exemplified by marker genes with recognized cell-type specificity, including *DCT2* (bundle sheath), *CARBONIC ANHYDRASE1* (*CAH1*; mesophyll), *POTASSIUM CHANNEL 3 (KCH3;* guard cell), *GLYCEROL-3-PHOSPHATE ACYLTRANSFERASE 12* (*GPAT12*; epidermis), and homolog to *GLABRA3* (*ZmGL3*; trichome) (**Figure 2E** and **2F**). Comparison of aggregated cell-type profiles indicated a greater extent of variation in chromatin accessibility relative to gene expression, suggesting chromatin structure provides additional information for dissecting cell-type heterogeneity (**Figure 2G**).

**Figure 2.**
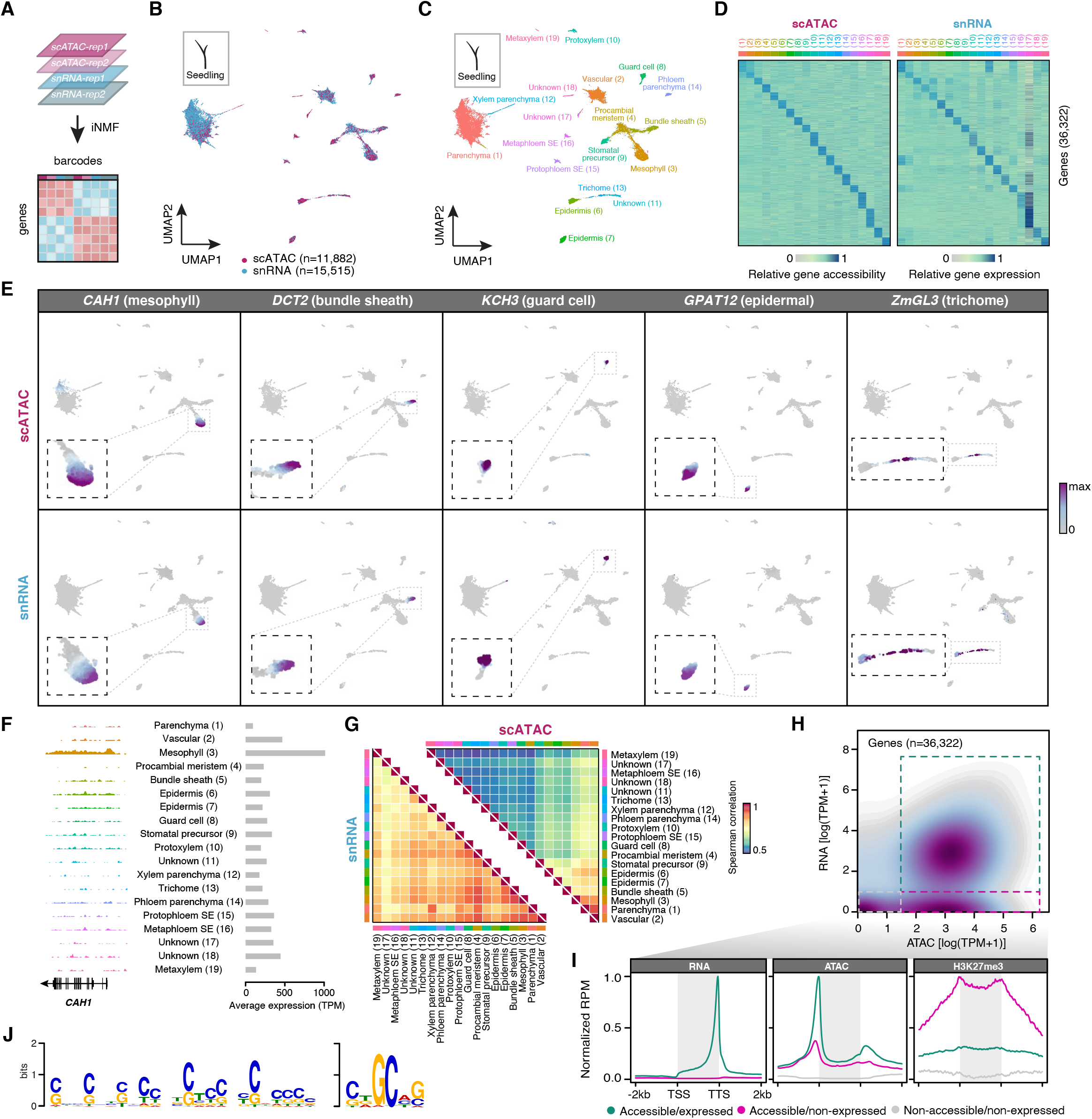
Concordance between chromatin accessibility and gene expression at single-nuclei resolution. (A) Illustration of iNMF integration of scATAC-seq and snRNA-seq seedling data sets. (B) UMAP co-embedding of seedling nuclei from scATAC-seq (n=11,882; purple) and snRNA-seq (n=15,515; blue). (C) Louvain clustering and cell-type annotations for co-embedded seedling nuclei. (D) Comparison of within-cluster averaged gene accessibility (left) and gene expression (right) between cell types/clusters. Column color legend corresponds to the cell-type colors specified in panel C. (E) UMAP embeddings displaying per nucleus gene accessibility (top, n=11,882) and gene expression (bottom, n=15,515) values for five cell type-specific marker genes. (F) Left: Aggregate scATAC-seq tracks across clusters at the *CAH1* locus. Right: Average expression of *CAH1*. (G) Spearman correlations between clusters based on nuclear transcription (snRNA) and chromatin accessibility (scATAC). (H) Density scatter plot comparing gene accessibility (x-axis) and expression (y-axis) for each cluster and gene. (I) Expression (left), chromatin accessibility (middle), and H3K27me3 ChIP-seq meta-profiles (relative reads per million, RPM) of accessible/expressed genes (turquoise; n=19,402), accessible/non-expressed genes (pink; n=6,063) and non-accessible/non-expressed genes (grey, n=4,315). (J) Top two *de novo* motifs enriched in ACRs within 1-kb of accessible/non-expressed genes (pink, panel H and I).

Despite the strong association between gene accessibility and expression, we observed a subset of accessible genes lacking evidence of transcription that were highly enriched with H3K27me3 (**Figure 2H** and **2I**). As we defined gene accessibility to include upstream sequences, we posited that ACRs associated with accessible and silenced genes might contain Polycomb Response Elements (PREs) directing transcriptional silencing via deposition of H3K27me3 by the Polycomb Repression Complex (PRC). *De novo* motif analysis of 15,073 ACRs within 1-kb of accessible and silenced genes (n=6,063) identified several enriched motifs, including a CNN-repeat (E-value < 2.0e-738, 83% of ACRs, 12,858/15,073) and a CTGCAG palindromic motif (E-value < 2.4e-205, 80% of ACRs, 12,014/15,703) (**Figure 2J**, **STAR Methods**). A query with experimentally established TF binding sites revealed a significant (FDR < 4.09e-3) overlap between the CNN-repeat motif and sequences recognized by BASIC PENTACYSTEINE1 (BPC1), a BARLEY B RECOMBINANT-BASIC PENTACYSTEINE (BBR-BPC) family TF previously associated with PREs and H3K27me3-mediated silencing in *A. thaliana* (Xiao et al., 2017) (**STAR Methods**). Taken together, we establish gene accessibility as a robust proxy for transcription and suggest the activity of PREs as a possible explanation for imperfect correlations between gene accessibility and expression.

To comprehensively investigate the extent of gene accessibility variation, we performed differentially accessibility hypothesis testing for each gene model across cell types (**STAR Methods**). After multiple test correction and heuristic thresholding (FDR < 0.05, log fold-change > 2), we identified 74% (28,625/38,752) of genes with significant differential accessibility in at least one cell type, with an average of 2,768 differentially accessible genes per cluster (**Figure S6A**; **Table S5**). Marker-agnostic gene set enrichment analysis (GSEA) of Gene Ontology (GO) terms exemplified prior information regarding specific cell-type functions, with enriched terms such as “root hair cell development” in root epidermal initials, “regulation of stomatal closure” in subsidiary cells, and “malate transmembrane transport” for mesophyll cells (**Figure S6D**). Distinct cell types were generalized by highly specific GO annotations, as most (>51%) GO terms were identified in only a handful of cell types (five or fewer), implicating chromatin accessibility dynamics as underling the signature hallmarks of cell-type identity and function (**Figure S6D**). In summary, we identified 52 cell types for 83% (76/92) of scATAC-seq clusters, capturing nearly all major expected cell types in the profiled organs and suggesting the existence of novel uncharacterized cell types present in these data (**Table S4**).

### Characterization of *cis*-regulatory variation

Deconvolution of nuclei into discrete cell types provides an opportunity to identify CREs encoding cell identity. To this end, we implemented a generalized linear model to catalog ACRs with discrete patterns of chromatin accessibility across cell types (**STAR Methods**). In total, 52,520 ACRs (31%) were differentially accessible (FDR < 0.05, fold-change > 2) and restricted to one or a handful of clusters, with an average of 2,826 per-cluster (**Figure S6B**, **S7A**, and **S7B**). Similar to the reported functions of transcriptional enhancers in metazoan genomes, we found cluster-specific ACRs were associated with significantly greater enhancer activity based on Self-Transcribing Active Regulatory Region sequencing (STARR-seq) of maize leaf protoplasts (Ricci et al., 2019) relative to controls (n=165,913) and non-specific ACRs (n=113,393; **Figure 3A**; **STAR Methods**). Deconvolution of chromatin accessibility by cell type revealed accessible sites primarily located distal to genic regions (>2-kb from any gene) compared to previously published bulk-level experiments (**Figure 3B**) (Ricci et al., 2019). Notably, 30% (22,456/73,791) of distal ACRs overlapped with LTR transposons, including the major maize domestication locus *TEOSINTE-BRANCHED 1-*enhancer (*tb1-*enhancer), and were generally devoid of DNA methylation (**Figure 3B**, **3C**, and **S1**) (Crisp et al., 2020; Oka et al., 2020; Oka et al., 2017). These findings are consistent with transposable elements playing a prominent role in CRE evolution of the maize genome (Clark et al., 2006; Noshay et al., 2020; Zhao et al., 2018).

**Figure 3.**
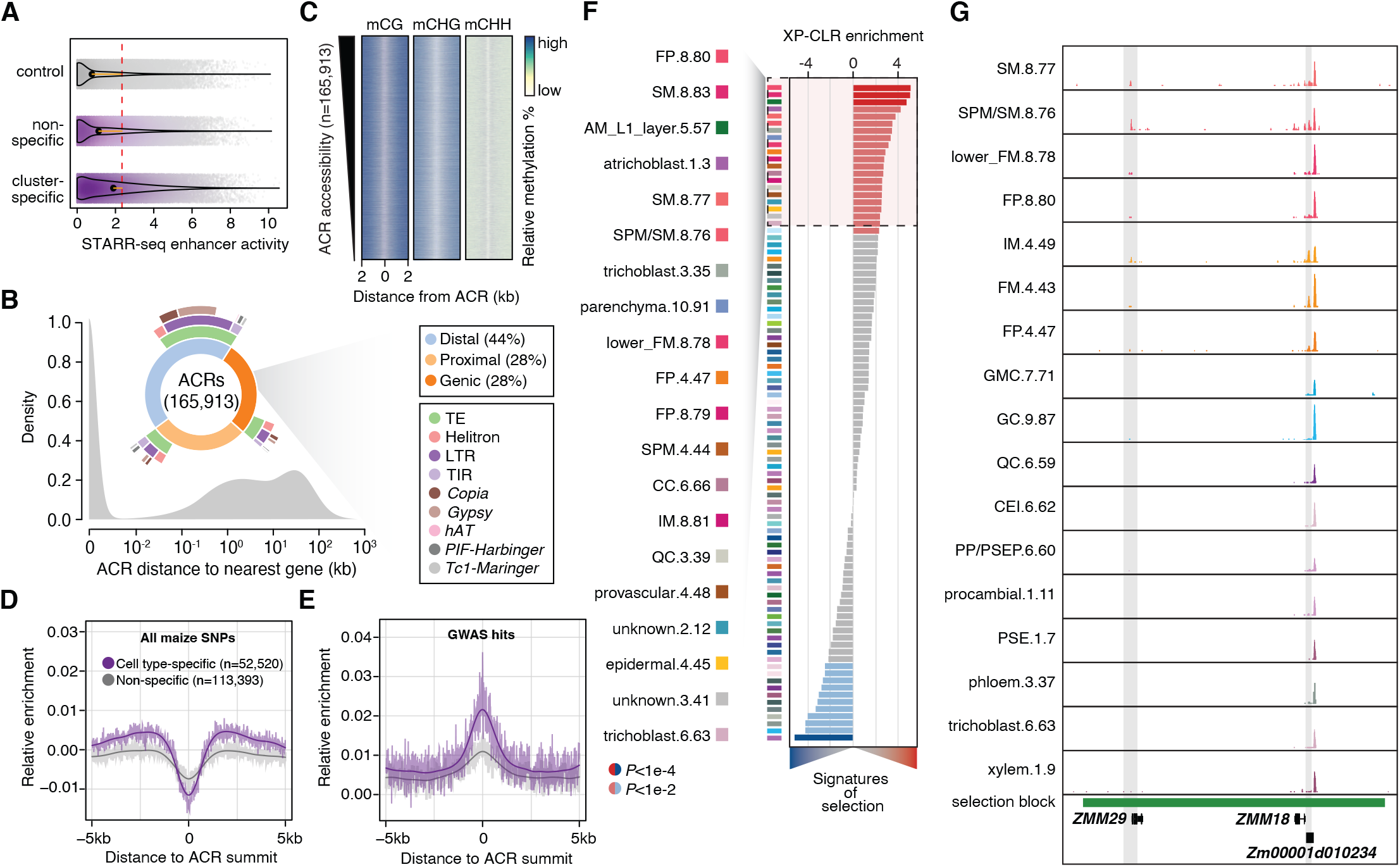
Characterization of accessible chromatin regions. (A) Distribution of enhancer activity (maximum log2[RNA/input]) for control regions (n=165,913), non-specific (n=113,393) and cluster-specific ACRs (n=52,520). The dash red line indicates the overall mean. Orange lines reflect differences between the group median and overall mean. (B) Distribution of ACR distances to the nearest gene. Inset, distribution of ACR genomic context. (C) Relative DNA methylation levels 2-kb flanking ACRs. (D) Relative enrichment of polymorphisms after normalizing by mappability for 5-kb regions flanking cell type-specific and non-specific ACR summits. Smoothed splines are shown as dark lines. (E) Relative enrichment of significant GWAS polymorphisms relative to all polymorphisms for 5-kb regions flanking cell type-specific and non-specific ACR summits. (F) Enrichment of signatures of selection (XP-CLR) in the top 2,000 ACRs for all cell-type clusters. The 20 most enriched cell types are highlighted on the left. AM, axillary meristem; CC, companion cell; FM, floral meristem; FP, floral primordia; IM, inflorescence meristem; QC, quiescent center; SM, spikelet meristem; SPM, spikelet pair meristem. (G) Aggregate scATAC-seq tracks for seven floral cell types and a random assortment of 10 non-floral cell types at *ZMM29* and *ZMM18* loci. CEI, cortex/endodermis initials; GC, guard cell; GMC, guard mother cell; PP/PSEP, pre-procambial/phloem sieve element precursor.

Sequence variation underlying CREs contribute to disease emergence and phenotypic innovation over evolutionary timescales (Rebeiz and Tsiantis, 2017; Villar et al., 2015). In contrast to broadly accessible chromatin regions, analysis of extant genetic variation in maize revealed lower polymorphism rates within cell type-specific ACRs (**Figure 3D**). However, of the genetic variants embedded within ACRs, those within cell type-specific ACRs were more frequently associated with phenotypic variation (Wallace et al., 2014) (**Figure 3E**). To investigate the contribution of domestication and selection in distinct cell-type contexts, we assessed the relative enrichment of selection signatures from chronologically sampled elite inbred maize lines within cell type-specific ACRs (**STAR Methods**) (Wang et al., 2020). Of the 21 cell types with significant (FDR < 0.01) selection signature enrichment, 57% (12) correspond to staminate and pistillate cell types, such as spikelet meristems, spikelet pair meristems, inflorescence meristems, floral meristems, and floral primordia (**Figure 3F**). For example, a single block encompassing two adjacent class B floral-organ morphology loci, *ZEA MAYS MADS 29* (*ZMM29*) and *ZMM18*, exhibited inflorescence, spikelet, and floral meristem and primordia-specific ACRs at both TSSs (**Figure 3G**). These findings indicate that modern maize breeding resulted in the selection of alleles containing floral-specific ACRs associated with agronomically favorable inflorescence architecture (Gage et al., 2018).

### Variation in transcription factor activities underlies cell identity

Differential TF binding has been proposed as a key driver of differential gene expression signatures underlying diverse cell identities. In line with recognized *cis*-regulatory function, ACRs were highly enriched with putative TF binding sites relative to control (n=165,913) and flanking regions, and were strongly depleted within ACR summits, consistent with TF-bound sequence occluding Tn5 integration (**Figure 4A**). Using the top 2,000 differential ACRs for each cell type, we found 84% of TF motifs (475/568) were enriched (binomial test: FDR < 0.05; **STAR Methods**) in at least one cell type, with a median of 43 enriched TF motifs per cell type (**Figure 4B**).

**Figure 4.**
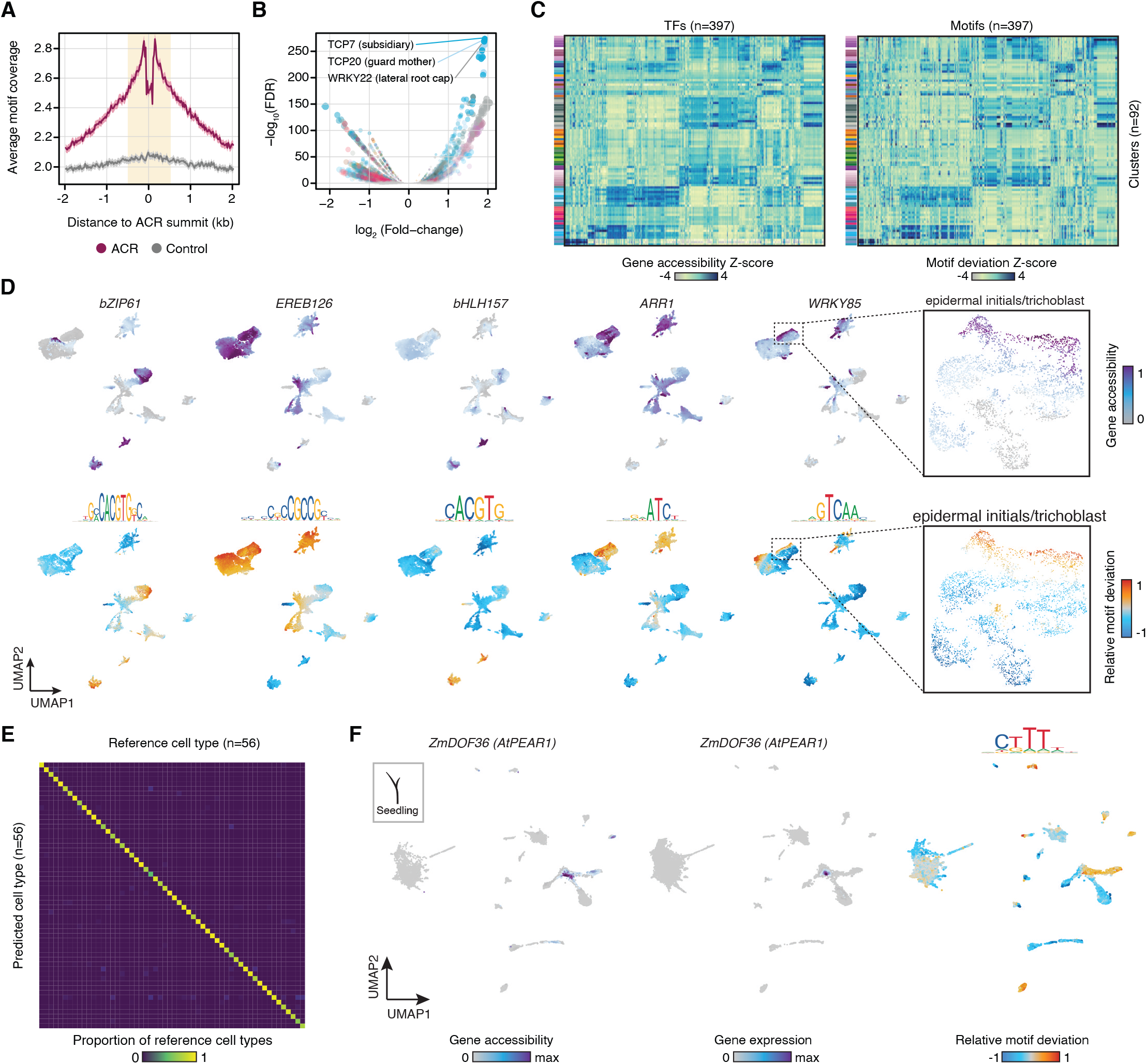
Combinatorial accessibility of transcription factor motifs and genes contribute to distinct cell identities. (A) Average motif coverage for all ACRs (n=165,913) and control regions (n=165,913). Shaded polygon, 95% confidence intervals. (B) Enrichment of TF motifs in the top 2,000 ACRs ranked by Z-score for each cell type compared to the top 2,000 most constitutive ACRs via binomial tests. FDR was estimated by the Benjamini-Hochberg method. (C) Comparative heatmaps for matched TF gene accessibility (bottom) and motif deviation (top) Z-scores across clusters. (D) Gene accessibility scores for five maize transcription factors (top) and their associated motif deviations (bottom). (E) Comparison of predicted vs. reference cell-type annotations from a neural-network multinomial classifier trained on combinatorial motif deviation scores. (F) Co-embedded seedling nuclei gene accessibility, RNA expression, and motif accessibility for *ZmDOF36*.

Next, we hypothesized that the relative accessibilities of motif across ACRs in a single nucleus could be used to elucidate the regulatory rules governing discrete cell states (**Figure S7C–S7F**). Comparison of TF gene accessibility with the relative accessibility of their sequence-specific binding sites revealed strikingly similar patterns across cell types (median Pearson’s correlation coefficient across cell types = 0.45), establishing synchronized chromatin accessibility of *cis* and *trans* cell-autonomous factors as major determinants of cell identity (**Figure 4C**, **4D**, and **S7G**). Assessment of enriched TFs and their cognate motifs identified several known cell type-specific regulators – including *WRKY* family TFs in root epidermal progenitors and trichoblasts (Verweij et al., 2016), *G2-like1* in parenchymal mesophyll (Chang et al., 2012), and *AGAMOUS*-*like* and *SEPALLATA* (Gomez-Mena et al., 2005) TFs in floral primordia – as well as previously uncharacterized TFs with new potential roles as cell-type regulators (**Figure 4C** and **4D**; **Table S6** and **S7**). To determine the utility of TF motif signatures for discerning cell identity, we trained a neural network (NN) on patterns of TF motif accessibility underlying various cell types. The NN model achieved an overall accuracy of 0.94 and an average sensitivity and specificity of 0.93 and 0.99, respectively, indicating that patterns of motif accessibility enable highly predictive classifications of diverse cell states (**Figure 4E**).

Past developmental genetic studies have described a handful of mobile TFs capable of influencing the identities of neighboring cells. As a proxy for non-cell autonomous activity, we searched for TFs with increased motif accessibility in cell types lacking expression (and accessibility) of the cognate TF. Of 279 TFs, we identified 20 with putative non-autonomous activity, including at least four TFs, *PHLOEM EARLY DOF1 (PEAR1), TEOSINTE BRANCHED1/CYCLOIDEA/PROLIFERATING CELL NUCLEAR ANTIGEN FACTOR4* (*TCP4*), *TCP5, TCP14*, and *ETHYLENE RESPONSE FACTOR 018* (*ERF018*), with predicted or known cell-cell mobility (Miyashima et al., 2019; Nag et al., 2009; Savaldi-Goldstein et al., 2007; Tatematsu et al., 2008). For example, *PEAR1* was recently described as a mobile DOF TF expressed in the procambium functioning to promote radial growth in the vasculature of *A. thaliana*. Consistent with predicted mobility, the maize *PEAR1* homolog, *ZmDOF36*, was largely expressed in procambial and protophloem cells, while its target motif was enriched in procambial, bundle sheath, phloem parenchyma, meta/protophloem, xylem, and epidermal cell types (**Figure 4F**; **Table S8**). These results indicate robust inference of CRE and TF activity at the level of single nuclei and reveal TF dynamics central to *cis*-regulatory specification of diverse cell states.

### Coordinated dynamic chromatin accessibility recapitulates *in vivo* chromatin interactions

Correlated changes in chromatin accessibility of nearby loci represent putative physical chromatin interactions with regulatory potential (Buenrostro et al., 2015; Cusanovich et al., 2018; Gate et al., 2018; Pliner et al., 2018; Satpathy et al., 2019). We identified 3.8 million (M) ACR-ACR linkages (hereafter referred to as co-accessible ACRs) with significantly correlated patterns of chromatin accessibility across cell types, capturing known gene-to-CRE physical interactions for gene loci such as *tb1,* maize *RELATED TO AP2.7* (*ZmRAP2.7*), and *BENZOXAZINLESS 1* (*BX1*) (empirical FDR < 0.05; **Figure 5A**, **S8**, **S9A**; **STAR Methods**) (Clark et al., 2006; Peng et al., 2019; Ricci et al., 2019; Salvi et al., 2007; Sun et al., 2020; Zheng et al., 2015). To assess the broad interactive potential of co-accessible ACRs *in vivo*, we compared co-accessible ACRs from seedling cell types with maize seedling chromatin conformation capture data, recovering more than 78% (3,313/4,265), 57% (37,712/65,691) and 44% (17,108/38,567) of chromatin loops from Hi-C, H3K4me3-HiChIP and H3K27me3-HiChIP experiments, respectively (**Figure S9B**). Hi-C/HiChIP is a direct reflection of the proportion of cells exhibiting a particular interaction, as ubiquitously interacting loci dominate rarer cell type-specific contexts that are frequently missed by loop-calling algorithms. To determine the relative predictability of *in vivo* interactions using co-accessible ACRs, we estimated the interaction strength of each ACR by integrating correlative scores across cell types (**STAR Methods**). ACRs classified by interaction strength recapitulated expected *in vivo* chromatin interaction frequencies, where even the weakest class of co-accessible ACRs were associated with elevated interaction frequencies relative to flanking and randomized control regions (**Figure 5B**).

**Figure 5.**
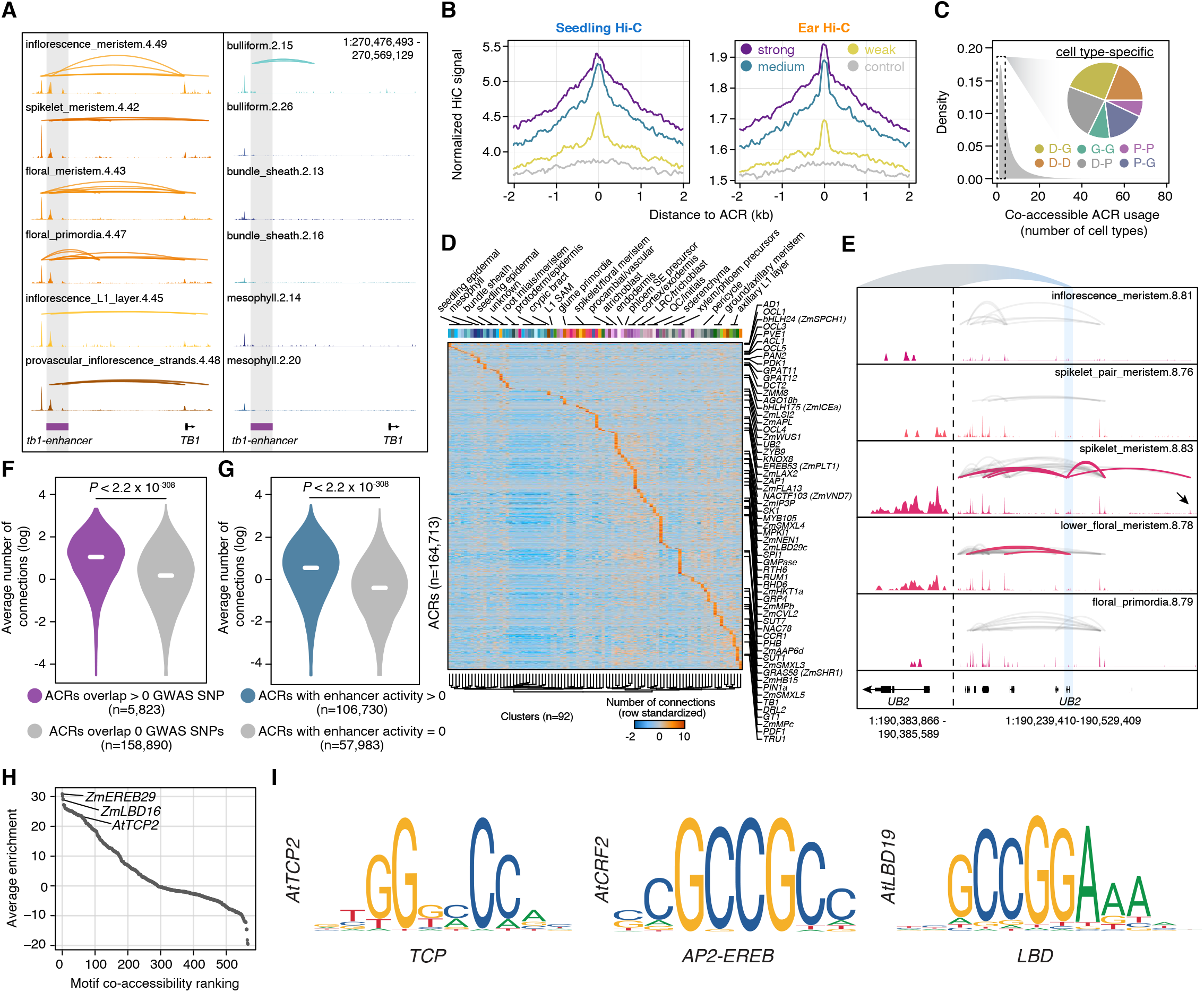
Co-accessible ACRs reflect *in vivo* chromatin interactions and are established by co-accessible TFs. (A) Comparison of ear and seedling co-accessible ACRs across nine cell types at the genetically mapped *tb1-enhancer* domestication locus (highlighted region). Link height reflects co-accessibility scores between ACRs across cells in a cluster. (B) Average normalized Hi-C signal across 4-kb windows centered on ACRs equally distributed into three groups (each group: n=54,904) based on the number and strength of participating co-accessible links. (C) Co-accessible ACR interaction frequency across clusters. Inset: Single cell-type co-accessible links (n=1,018,417) for different genomic contexts. D, distal; P, proximal; G, genic. Example: D-P indicates co-accessible ACRs where one edge is distal (> 2-kb from any gene) and the other is proximal (< 2-kb from any gene). (D) Standardized (row-wise) number of connections for each ACR (rows) by cell type (columns). Column color map reflect cell types from the legend in Figure S6. Gene-proximal ACRs for a subset of marker genes are indicated on the right. (E) Right, chromatin accessibility and co-accessible ACR links surrounding the *UB2* locus associated with ear row number and tassel branch number quantitative traits. Black arrow indicates a distal ACR upstream of *UB2* present only in spikelet meristems. Co-accessible links with an edge within 2-kb of *UB2* are colored pink while remaining links are grey. Link height represents co-accessibility strength. Left, close-up of accessibility profiles of *UB2*. (F) Distributions of average ACR-ACR links across cell types for ACRs that overlap (purple) and do not overlap (grey) phenotype-associated genetic variants from maize GWAS. The median of each distribution is shown as a white horizontal line. Violin plots present the entire range of average number of connections on a log scale. Hypothesis testing was conducted within the *R* statistical framework via Wilcoxon rank sum test. (G) Distributions of average ACR-ACR links across cell types for ACRs with (blue) and without (grey) enhancer activity (log_2_ RNA/input greater than 0). Hypothesis testing and distribution illustration was performed similarly as panel F. (H) Motifs ranked by the average co-accessibility enrichment over background across all cell types. (I) Exemplary motifs enriched in reciprocal co-accessible ACRs for *TCP*, *AP2-EREB*, and *LBD* TF families.

Cataloguing the usage of co-accessible ACRs across cell types identified more than 27% (∼1M in total) that were unique to a single type, 49% of which were classified as distal-genic or distal-proximal, and an average of 11,069 distinct links per cell type (**Figure 5C**). Consistent with regulatory models where a single gene can interact with multiple distant loci, proximal ACRs, rather than distal or genic ACRs, were associated with the greatest number of links on average (Wilcoxon rank sum test: *P* < 2.2e-308; **Figure S9C**). Highlighting long-range “hub” interactions as key contributors towards cell identity, cell type-specific co-accessible ACRs were associated with greater number of links per site (Wilcoxon rank sum test: *P* < 2.2e-308) and a greater proportion of links involving distal ACRs (Chi-squared test: *P* < 2.2e-308; **Figure S9C** and **S9D**). Furthermore, the interactive capacity of any given ACR strongly depended on the cell-type context (**Figure 5D**). For example, *UNBRANCHED 2* (*UB2*) – a major ear row number and tassel branch number quantitative trait locus (Chuck et al., 2014) – demonstrated preferential accessibility in spikelet meristems that coincided with the greatest number of *UB2* proximal to distal ACR interactions, including a cell type-specific ACR located upstream approximately 150-kb (**Figure 5E**). We posited that ACRs with expanded interactive capacity resemble enhancers with the potential to influence organismal phenotypes. Indeed, ACRs with enhancer activity and co-localization with phenotype-associated genetic variants from GWAS were associated with a significantly greater number of ACR-ACR connections (Wilcoxon rank sum: *P* < 2.2e-308; **Figure 5F** and **5G**). These results highlight the occurrence of diverse cell type-specific regulatory configurations among distal enhancer ACRs and their target genes and implicate genetic variants perturbing highly interactive distal enhancers as major contributors towards phenotypic variation.

The structural protein, CCCTC-binding factor (CTCF) plays an important role in metazoan genome organization and is notably absent in plant lineages (Heger et al., 2012). In search of an orthogonal factor in maize, we hypothesized that higher-order chromatin structure captured by co-accessible ACRs may be driven by TFs recognizing similar sequence motifs embedded within interacting accessible regions. Comparison of motif occurrences between co-accessible link edges indicated that ACRs in co-accessible links are more similar to one another than randomly linked ACRs (empirical: *P* < 1e-4; **Figure S9E**). Furthermore, we identified several cases where co-accessible ACR edges were reciprocally enriched for the same TF motif in both cell type-specific and non-specific co-accessible links (FDR < 0.05; **Figure S9F**; **STAR Methods**). Ranking motifs by the average enriched across cell types, we identified *TCP*, *APETALA2*/*ETHYLENE-RESPONSIVE ELEMENT BINDING PROTEINS* (*AP2-EREBP*) and *LATERAL ORGAN BOUNDARIES DOMAIN* (*LBD*) motifs that were not only broadly associated with co-accessible edges, but also exhibited strikingly similar GC-rich palindromic binding sites (FDR < 0.05; **Figure 5H**, **5I** and **S9F**). A role in chromatin organization is supported by previous research demonstrating *TCP* motif overrepresentation in topologically associated domain-like (TAD) boundaries in *Oryza sativa* and *Marchantia polymorpha,* and the distal edges of chromatin loops in *Z. mays* (Karaaslan et al., 2020; Liu et al., 2017; Peng et al., 2019; Sun et al., 2020). Consistent with these past studies, our results implicate independently evolved TF families with CTCF-like function capable of organizing higher-order chromatin architecture through DNA-protein interactions.

### Dynamic chromatin accessibility specifies cell developmental trajectories

The apical domains of maize enclose a pool undifferentiated meristematic stem cells that give continuous rise to all differentiated cell types (Somssich et al., 2016; Takacs et al., 2012). To define a *cis*-regulatory catalog of temporal cell fate progressions, we ordered nuclei along pseudo-temporal trajectories for 18 developmental continuums, reflecting meristematic to differentiated cell states; identifying ACRs, TF loci, and TF motifs with significant variation across pseudotime (**Figure 6A** and **S10**; **Table S9-S12**; **STAR Methods**). To showcase the power of trajectory construction to characterize a relatively understudied process, we focused our analysis on root phloem companion cell (PCC) development (**Figure 6B**). We identified 8,004 ACRs, 440 TF motifs, 7,955 genes, and 402 TF loci that were differentially accessible across the PCC pseudotime trajectory (**Figure 6C**; **STAR Methods**). Several known meristem and phloem developmental genes including *AT-RICH INTERACTIVE DOMAIN-CONTAINING 8* (*ARID8*) (Jiang et al., 2010), *SUPPRESSOR OF MAX2 1-LIKE3* (*ZmSMXL3*) (Wallner et al., 2017), *and SUCROSE TRANSPORTER 1* (*ZmSUT1*) (Baker et al., 2016), were identified among the top differentially accessible genes throughout PCC development (**Figure 6D**).

**Figure 6.**
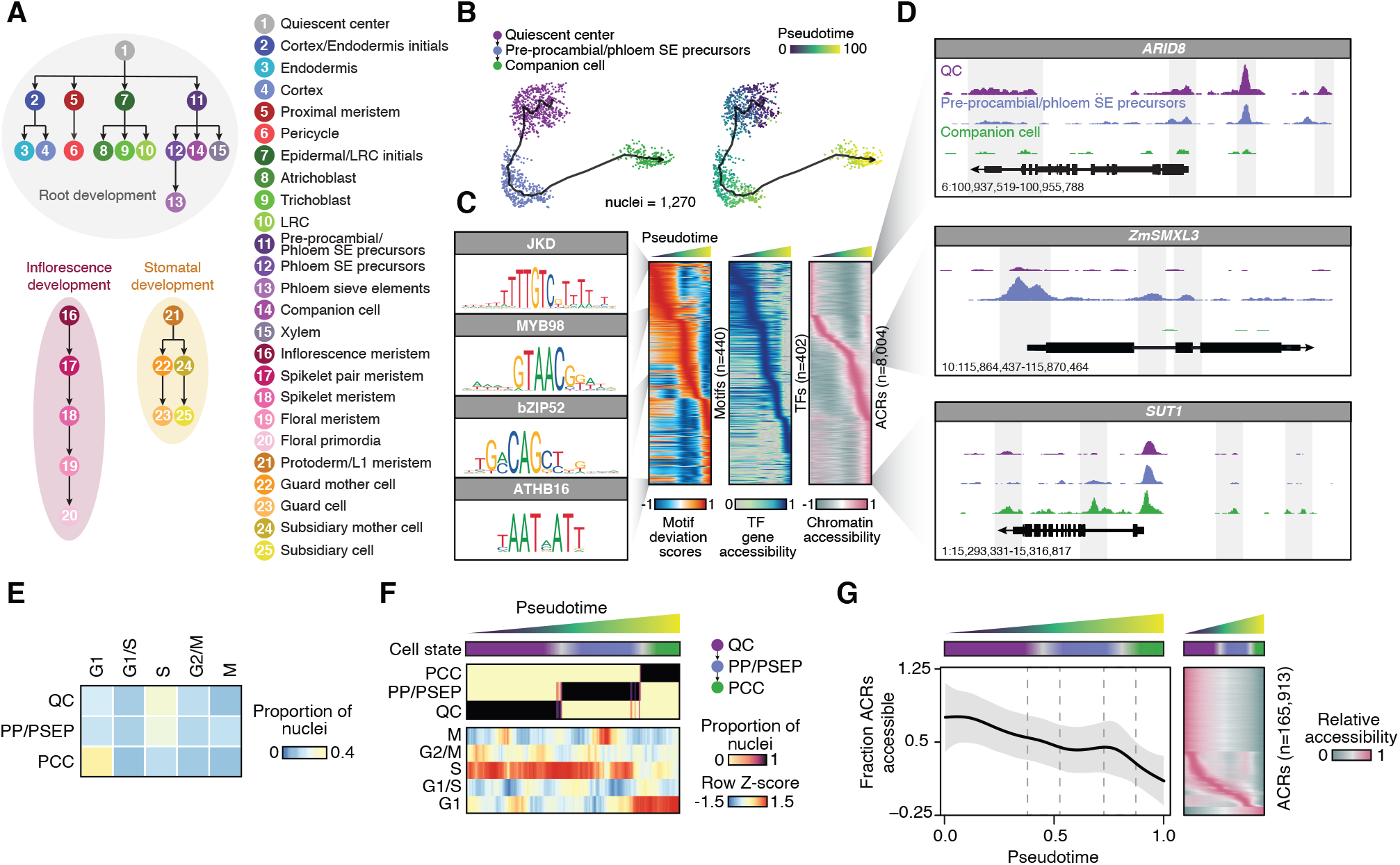
Chromatin accessibility is dynamic across pseudotime. (A) Overview of pseudotime trajectory analysis. Inflorescence development was performed for both pistillate and staminate inflorescence. (B) UMAP embedding of companion cell developmental trajectory depicting cell types (left) and pseudotime progression (right). (C) Relative motif deviations for 440 TF motifs (left, rows), 402 TF gene accessibility scores (middle, rows), and relative accessibility of 8,094 ACRs (right, rows) associated with pseudotime (columns). Four motifs enriched along the trajectory gradient are shown on the left. ACRs, accessible chromatin regions; TF, transcription factor. (D) Genome browser screenshots of cell type-specific chromatin accessibility profiles along the developmental trajectory for quiescent center (QC), pre-procambial/phloem sieve element precursor (PP/PSEP), and phloem companion cell (PCC) at associated marker gene loci. (E) Proportion of cells at various stages of the cell-cycle in QC, PP/PSEP, and PCC annotated clusters. (F) Top: Cell state ordered by pseudotime. Middle: Proportion of nuclei with the corresponding cell-type annotation ordered by pseudotime. Bottom: proportion of nuclei with various cell-cycle stage annotations ordered by pseudotime. Nuclei were binned into 250 blocks. (G) Left: Average fraction of ACRs that are accessible across pseudotime. The grey polygon indicates standard deviation. Red windows indicate cell state transitions. Right: heatmap of relative accessibility (relative to the row maximum) for each ACR (rows) across pseudotime (columns). Nuclei were binned into 250 blocks ordered on pseudotime.

Past studies of root cell fate decisions have focused on the role of cell cycle in establishing patterns of asymmetric cell division, with quiescent center/meristematic cells dividing much slower than cells in the rapidly dividing transition and elongation zones (Ten Hove and Heidstra, 2008). To investigate the contribution of cell-cycling to PCC development, we annotated nuclei using *a priori* compiled list of known cell-cycle marker genes (**STAR Methods**) (Nelms and Walbot, 2019). As consequence of slower DNA replication and consistent with previous reports, the majority of QC and meristem/initial-like nuclei were in S-phase, while differentiated companion cells largely presented as G1 (**Figure 6E**). Ordering nuclei by PCC pseudotime indicated sequential progression of cycle stages within each cell type, revealing the cell cycle context preceding cell fate transitions along the PCC trajectory (**Figure 6F**). Furthermore, evaluation of global accessibility across pseudotime illustrated steady decrease in chromatin accessibility throughout PCC development (**Figure 6G**). Thus, cell-cycling and cell fate transitions in the context of PCC development accompany global decreases in chromatin accessibility, a consequence we posit is associated with acquisition of more specialized functions in PCCs relative to their meristematic progenitors.

### Evolutionary innovation in root development

Despite nearly 150M years of divergence, monocot and eudicot angiosperm species exhibit remarkable phenotypic similarity with core organs such as seeds, roots, and shoots functionally maintained. To explore the degree of regulatory conservation in angiosperm root development, we profiled chromatin accessibility in 4,655 nuclei from 7-day old embryonic root tissues in the eudicot model species *A. thaliana*, integrated with previously generated *A. thaliana* root scRNA-seq data (n=12,606), and constructed eight *cis*-regulatory pseudotime trajectories encompassing vascular, dermal and ground development (**Figure 7A–7C**, **S11** and **S12A**; **Table S13-S17**).

**Figure 7.**
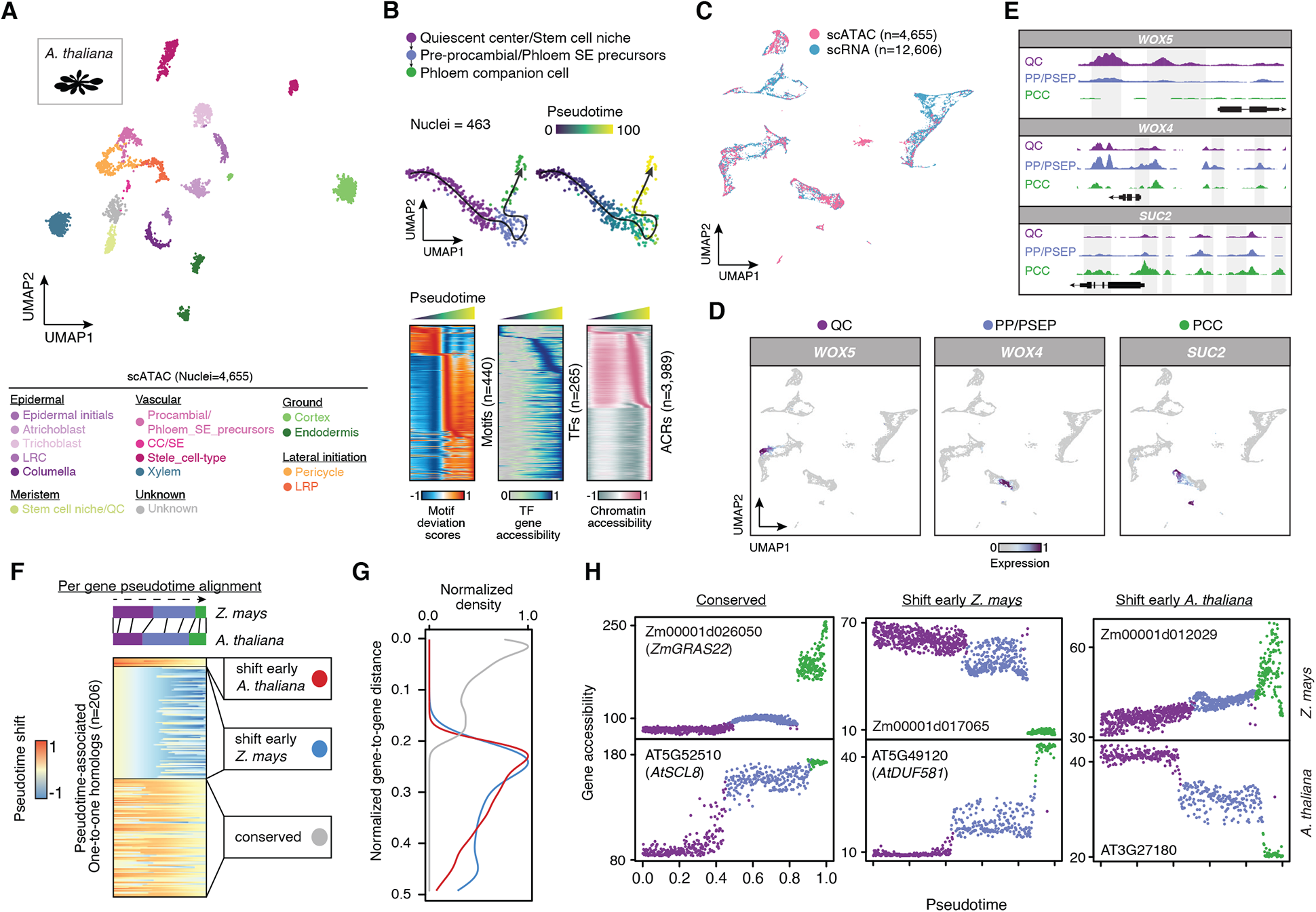
Dynamic and conserved *cis*-regulation in *A. thaliana* and *Z. mays* phloem companion cell development. (A) UMAP embedding based on whole-genome chromatin accessibility profiles of 4,655 *A. thaliana* root nuclei. (B) UMAP embedding of companion cell developmental trajectories in *A. thaliana* depicting cell types (left) and pseudotime progression (right). Motifs, TFs, and ACRs with significant association with pseudotime are shown as heatmaps below. (C) UMAP embedding of integrated scRNA-seq and scATAC-seq cell and nuclei profiles derived from *A. thaliana* roots. (D) Marker gene expression levels for individual scRNA-seq cells for quiescent center (QC; *WOX5*), pre-procambial/phloem SE precursors (PP/PSEP; *WOX4*), and phloem companion cells (PCC; *SUC2*). (E) Pseudobulk chromatin accessibility pile-ups for clusters labeled as QC, PP/PSEP, and PCC across three marker genes associated with each cell type, respectively. (F) Per-gene pseudotime shift scores from alignments between *Z. mays* and *A. thaliana* companion cell development progressions, clustered by k-means into three groups. (G) Distribution of gene-gene distances from the alignments, split by k-mean groups. (H) Exemplary one-to-one homologs between *A. thaliana* and *Z. mays* for the three groups split by pseudotime shifts. Acronyms: LRC, lateral root cap; LRP, lateral root primordia. SE, sieve elements.

Maintaining focus on PCC development, we first validated the utility of the integrated data sets by visualizing gene expression and accessibility of known marker genes representative of QC (*WUSCHEL-RELATED HOMEOBOX 5, WOX5*), procambial (*WUSCHEL-RELATED HOMEOBOX 4, WOX4*) and PCC (*SUCROSE TRANSPORTER 2, SUC2*) cell types (**Figure 7D** and **7E**). Next, we aligned *Z. mays* and *A. thaliana* PCC trajectories using a time-warping algorithm to enable direct comparison of gene accessibility dynamics in a common space. Consistent with recent comparative analysis of vascular development in *O. sativa*, *A. thaliana,* and *Solanum lycopersicum* (Kajala et al., 2020), only 206 out of 10,976 putative orthologs were significantly associated (FDR < 0.01) with PCC pseudotime in both species, indicating that the majority PCC trajectories-associated genes are unique to each lineage (97% *Z. mays*, 83% *A. thaliana*). However, of the 206 PCC-associated orthologs, ∼50% (102/206) exhibited similar patterns of gene accessibility across pseudotime (**Figure 7F**, **7G**, **S12B**). Several putative orthologs with matching gene accessibility patterns have been previously associated with PCC development, such as *SCARECROW-LIKE8* (*SCL8*), which displays an increasing expression gradient in nascent to mature *A. thaliana* PCCs in a previously generated *A. thaliana* root cell-type gene expression atlas (**Figure 7H**) (Brady et al., 2007). The remaining homologs (n=104) exhibited differential accessibility patterns clustered into two groups that reflect changes in the timing of gene accessibility across the PCC trajectory, underscoring putatively functional novelty in PCC development between *Z. mays* and *A. thaliana*.

To understand the extent of innovation in *cis* regulation along the pseudotime continuum, we aligned *Z. mays* and *A. thaliana* TF motif accessibility profiles associated with PCC progression (**Figure S12C**, **S12D**). Of the 440 motifs, 142 demonstrated highly conserved *cis*-regulatory dynamics between species (**Figure S12E–S12G**). Indeed, TFs recognizing the top four motifs ranked by normalized distances (**STAR Methods**) included *HOMEOBOX25* (*HB25*), *HOMEOBOX18* (*HB18*), *NAC DOMAIN CONTAINING PROTEIN 55* (*NAC055*), and *NAC DOMAIN CONTAINING PROTEIN 83* (*NAC083*) that have been previously implicated in regulation of hormonal responses and vascular development (Jiang et al., 2009; Yamaguchi et al., 2010; You et al., 2019). Gene expression profiles of these TFs from published root cell-type resolved data in *A. thaliana* was largely restricted to maturing procambial and companion cells, consistent with motif accessibility dynamics in both *Z. mays* and *A. thaliana* ontogenies (**Figure S12H**) (Brady et al., 2007). These finding signify a high degree of conservation in the *cis*-regulatory specification of PCC development between *Z. mays* and *A. thaliana* despite an analogous lack of concordance in accessibility dynamics among orthologous genes.

## DISCUSSION

Here, we describe 92 distinct cell-states from a suite of *Z. mays* organs, many of which previously lacked prior genome-wide characterization. Although we provide evidence supported by RNA *in situ* hybridization and snRNA-seq for the use of chromatin accessibility as a robust proxy of gene expression, the cell-type annotations should be considered preliminary. We anticipate that as single-cell methods become more widely adopted, including same-cell multi-modal experiments, these cell-type classifications, as well as those that were left unannotated, will become refined. An important consideration is that the sparse and binary nature of current scATAC-seq protocols require cell/nuclei aggregation by cell type for downstream analyses. Thus, additional gains in experimental procedures, particularly in reads per cell together with comprehensive transcriptome profiling within the same cell, will be necessary to fully investigate heterogeneity of cells within the same type. We anticipate that the application of same-cell multimodal techniques will open the door to better establish the molecular relationships among chromatin accessibility, gene expression, and cellular heterogeneity. Advancements in computational tools for comparing cell-type atlases and for data integration will play a key role in enabling higher resolution analyses than is currently possible.

Notwithstanding the technical challenges of single-cell experiments, our results represent a landmark advance for appreciating variation in cell-type functions established by diverse *cis*-regulatory grammar. We defined the TFs, CREs, and other loci that discretize cell-type identities and the sequential trajectories for an array of developmental ontogenies. Evaluation of motif accessibility variation alone was sufficient to predict cell identity with a high degree of accuracy, sensitivity and specificity. Querying patterns of motif accessibility relative to TF gene expression highlighted transcriptional regulators with putative non-cell autonomous activity, a particularly exciting result as identification of candidate non-cell autonomous factors previously relied exclusively on transgenic approaches to illuminate both transcript and protein localizations. Towards uncovering major regulators of global chromatin structure in species that lack CTCF, analysis of co-accessible ACRs successfully linked distal ACRs to their target genes and provided CTCF-like candidate TFs putatively orchestrating higher-order chromatin interactions that have been posited by orthogonal approaches. Further dissection of candidate regulatory genes and regions promises to be a fruitful endeavor for precise engineering of spatiotemporal patterns of gene expressions.

With an evolutionary perspective, we reveal that floral cell type-specific ACRs have been the historical targets of modern agronomic selection in maize. While subject to considerable sequence constraint, both by artificial and natural selection, genetic variation that does exist within cell type-specific ACRs is highly enriched with significant phenotypic variation. These findings point to abundant genetic variation capable of large phenotypic effects present within extant maize germplasm and present an ideal launchpad towards allele-mining for crop improvement. To understand the extent of cis-regulatory evolution in two highly diverged species, we constructed cell-type resolved chromatin accessibility profiles in the eudicot species *A. thaliana*. To our surprise, comparison of established orthologs indicated that a majority of genes involved in cell-type development were unique to each lineage. This finding contrasted with the observation of greater conservation of *cis*-regulatory elements involved in PCC development, as a greater proportion of TF motifs exhibited consistent spatiotemporal progressions among the two species. Viewed collectively, the maize *cis*-regulatory atlas presents an ideal framework for understanding the basis of cell heterogeneity, cis-regulatory control of gene transcription, and the foundation for future crop improvement efforts through targeted genome editing, synthetic biology approaches, and traditional allele-mining.

## AUTHOR CONTRIBUTIONS

APM and RJS designed the research. APM and ZC performed the experiments. APM, ZC, AG and RSJ analyzed the data. APM and RJS wrote the manuscript.

## ACKNOWLEDGEMENTS

We would like to acknowledge Dr. Zefu Lu for training in nuclei extraction, Dr. William Ricci for providing inflorescence samples, Julie Nelson from the University of Georgia flow cytometry core for helping sort nuclei, Noah Workman and Julia Portocarrero from the Georgia Genomics and Bioinformatics Core for library preparation, the Georgia Advanced Computing Resource Center for providing access to computing resources, and Tyler Earp for assistance with data management. This study was funded by support from the National Science Foundation (IOS-1856627 and IOS-1802848), the Pew Charitable Trusts and the UGA Office of Research to RJS, the National Science Foundation (IOS-1916804) to AG, and an NSF Postdoctoral Fellowship in Biology (DBI-1905869) to APM. AG and RJS are also supported by an NSF Collaborative award to support this research (IOS-2026554 and IOS-2026561). RJS is a Pew Scholar in the Biomedical Sciences, supported by The Pew Charitable Trusts.

## SUPPLEMENTAL FIGURE LEGENDS

**Figure S1:**
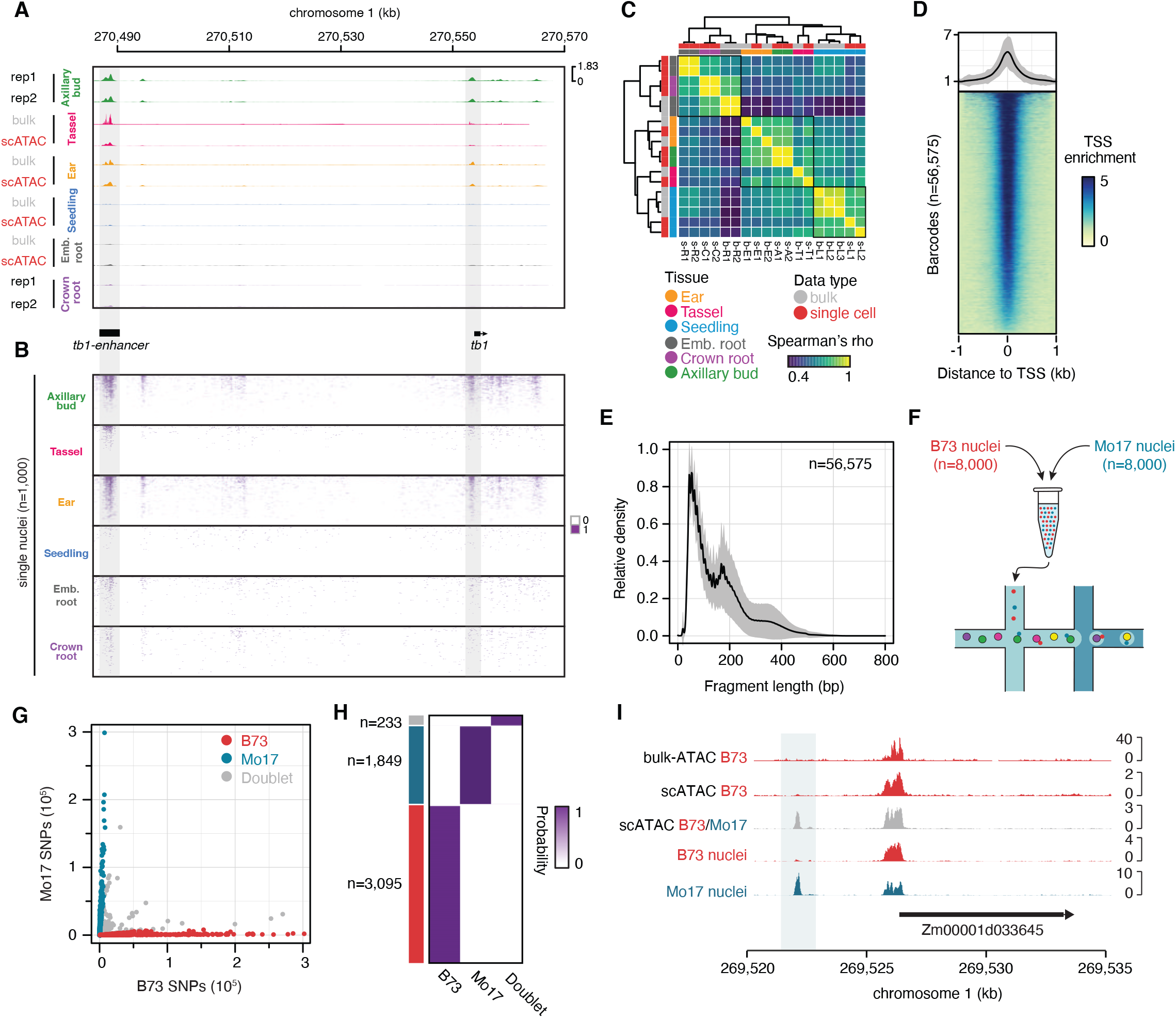
Evaluation and quality control of maize scATAC-seq. (A) Genome browser screenshot of chromatin accessibility from bulk and aggregated single-cell ATAC-seq experiments. Chromatin accessibility profiles depict the *tb1* locus and the *tb1* enhancer located approximately 67kb upstream. (B) Binary accessibility scores from a random selection of 1,000 individual nuclei from each organ. (C) Spearman’s rho matrix comparing bulk ATAC-seq and aggregate scATAC-seq samples across various organs. Sample codes are shorthand for assay type, sample, and replicate. For example, s-R1 denotes single cell assay for seminal root replicate 1. The term b-L2 denotes a bulk-ATAC assay for seedling replicate 2. Codes are as follows: b, bulk; s, single cell; R, seminal root; C, crown root; E, ear; T, tassel; A, axillary bud; L, seedling. Numbers represent replicate. (D) Average TSS enrichment (normalized read depth adjusted by the two 10 bp windows 1kb away from TSSs) across all 56,575 cells. Grey polygon denotes the standard deviation. (E) Fragment length distributions across 56,575 cells. The solid line and grey polygon represent the average and standard deviation, respectively. (F) Genotype-mixing experimental schematic. (G) Scatterplot of per cell B73 and Mo17 SNP counts from a mixed-genotype experiment (V1 seedlings) colored by genotype classification. (H) Posterior probabilities of individual barcodes (rows) highlighting the occurrence of cells with B73, Mo17, and mixed (doublet) genotype identities. (I) Genome browser screenshot of traditional bulk ATAC-seq from 7-day old seedling (row 1), single-cell ATAC-seq from B73 seven day old seedlings (row 2), pooled B73 and Mo17 nuclei (library ID: Seedling 2) single cell ATAC-seq from 7 day old seedlings (row 3), and the genotype-sorted B73 (row 4) and Mo17 (row 5) alignments after sorting barcodes by genotype calls from the B73-Mo17 scATAC-seq 7 day old seedling sample (row 3).

**Figure S2.**
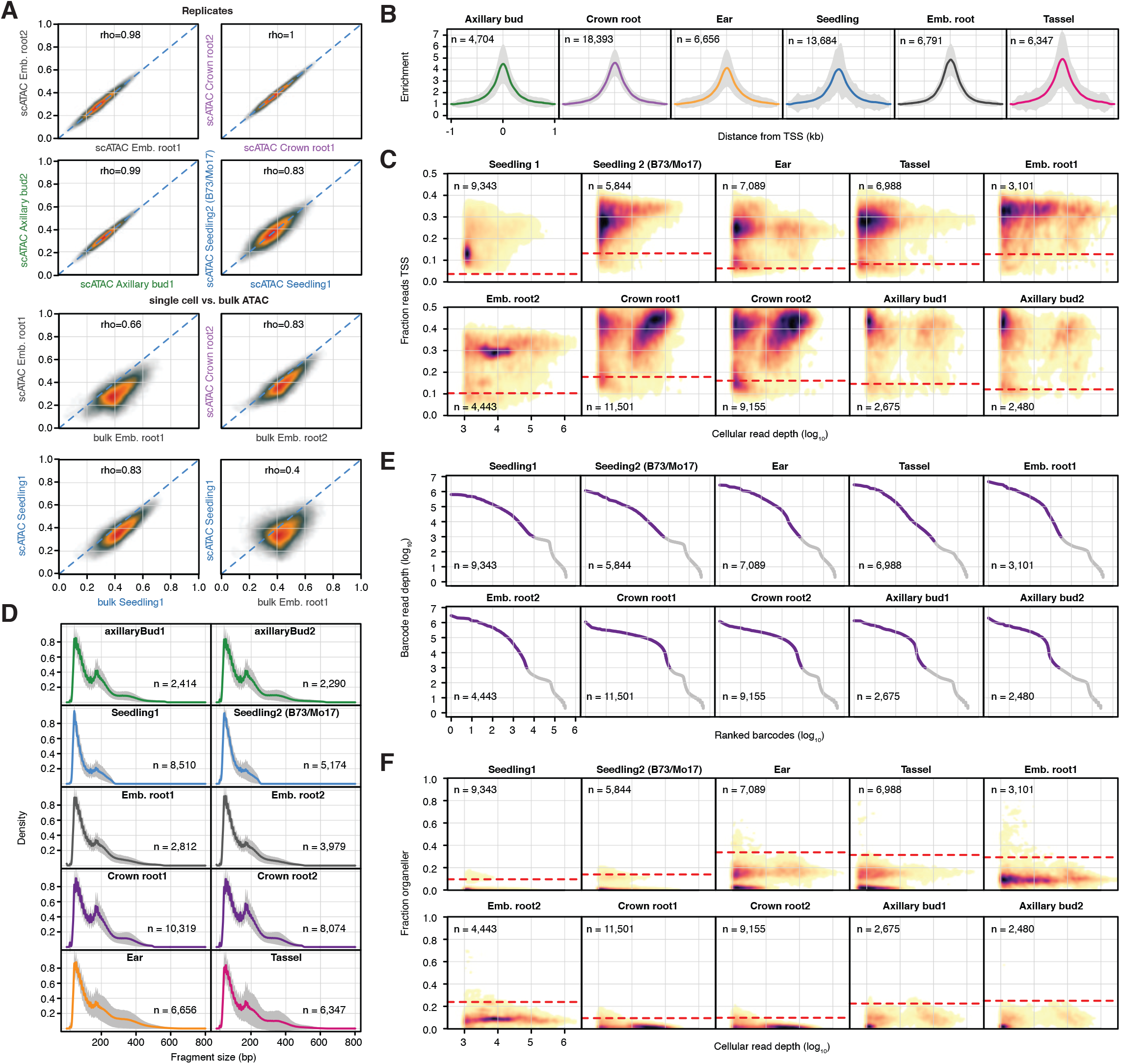
Cell calling and barcode quality control. (A) Comparison of normalized (0-1) read depths at the union of all peaks across bulk and single-cell samples (n=265,992) between replicated libraries, and between bulk and single-cell ATAC-seq assays. (B) Enrichment plots centered on 2-kb windows surrounding TSSs for barcodes in each tissue. Grey polygons indicate the standard deviation across cells within the noted tissue. (C) Density scatter plots of log_10_ transformed barcode read depths (x-axis) by the fraction of Tn5 integration sites mapping to within 2-kb of transcription start sites (TSSs). Dashed red lines indicate the threshold of two standard deviations from the mean used to filter lower quality barcodes. (D) Fragment length distributions for each library. Solid lines indicate the average distribution across cells within the sample. Grey polygons represent the standard deviation across cells in the library. (E) Knee plots illustrating log_10_ transformed cellular read depths of log_10_ ranked barcodes across libraries. (F) Density scatter plots of log_10_ transformed barcode read depths (x-axis) by the fraction of Tn5 integration sites derived from organeller sequences (chloroplast and mitochondrial) relative to the total number of unique Tn5 integration sites associated with cognate barcode. Dashed red lines indicate the threshold of two standard deviations from the mean used to filter lower quality barcodes.

**Figure S3.**
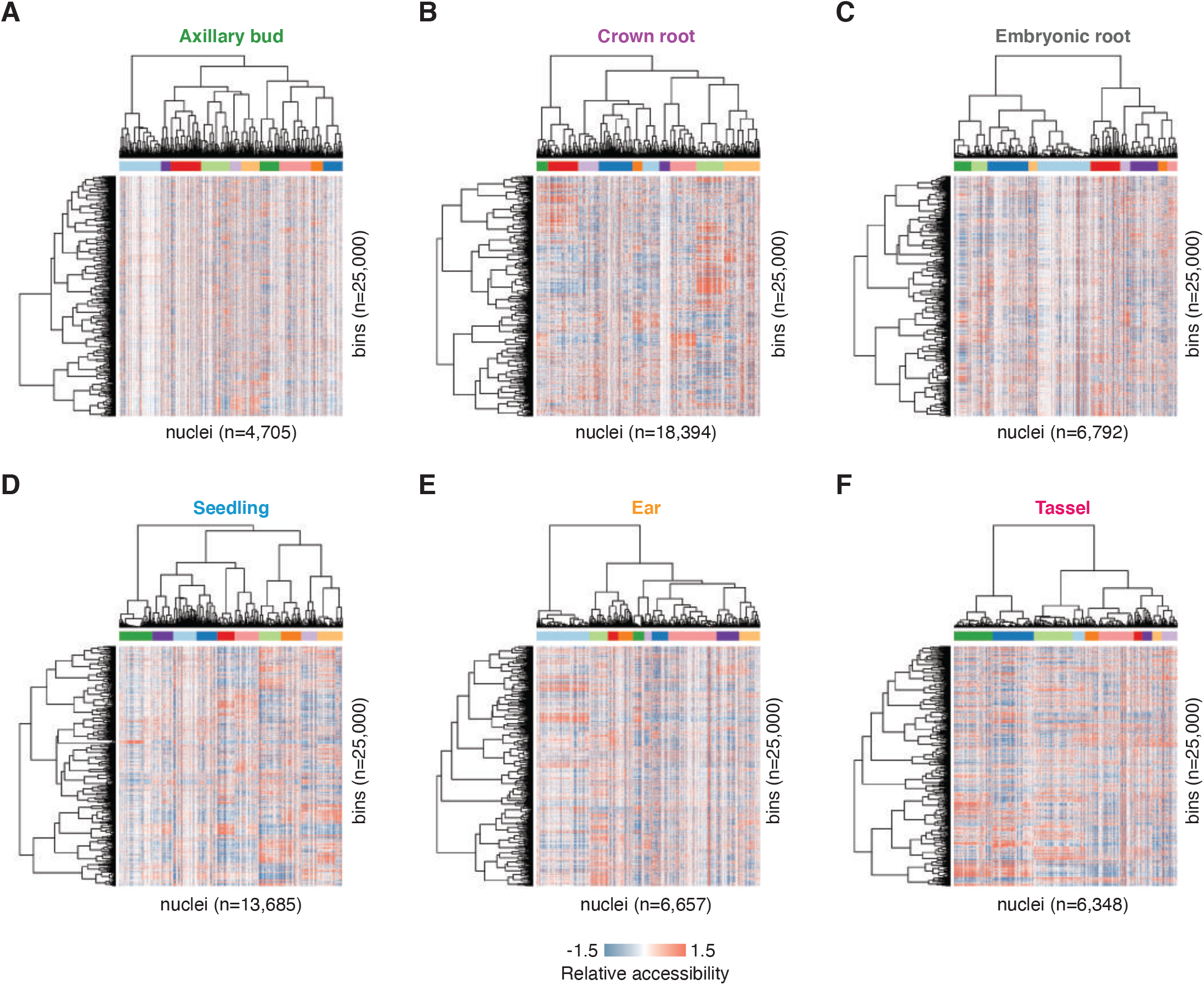
*In silico* sorting via Latent Semantic Indexing. Standardized LSI accessibility scores (2^nd^ - 11^th^ dimensions) capped at ±1.5 for: (A) Axillary buds. (B) Crown roots. (C) Embryonic roots. (D) Seedlings. (E) Ear (pistillate inflorescence). (F) Tassel (staminate inflorescence).

**Figure S4.**
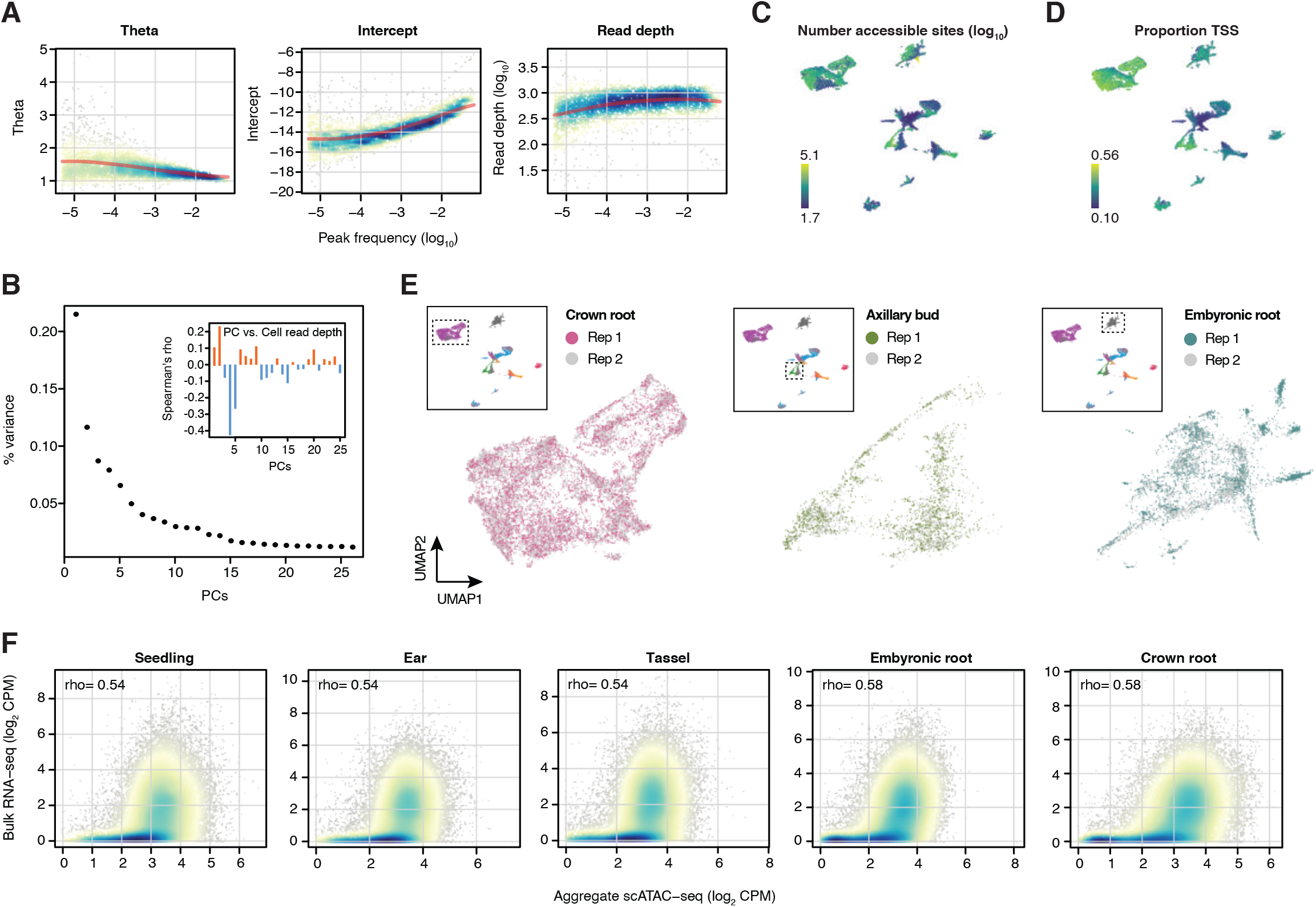
Clustering metrics and comparison of bulk gene accessibility and expression. (A) Parameter regularization of model coefficients (y-axes) with respect to ACR usage (x-axes; proportion of nuclei with at least one Tn5 integration site in an ACR). (B) Proportion of variance captured by the first 26 PCs. Inset: Spearman’s correlation of principal components with cell read depth (log_10_-transformed). (C) Number of accessible sites per cell (log_10_). (D) Proportion of Tn5 integrations within 2-kb of gene TSSs per cell. (E) Co-localization of nuclei barcodes from different biological replicates for three organs. (F) Comparison of bulk RNA-seq expression levels (y-axis, log_2_ CPM) versus aggregate scATAC-seq gene accessibility scores (x-axis, log_2_ CPM) within an organ.

**Figure S5.**
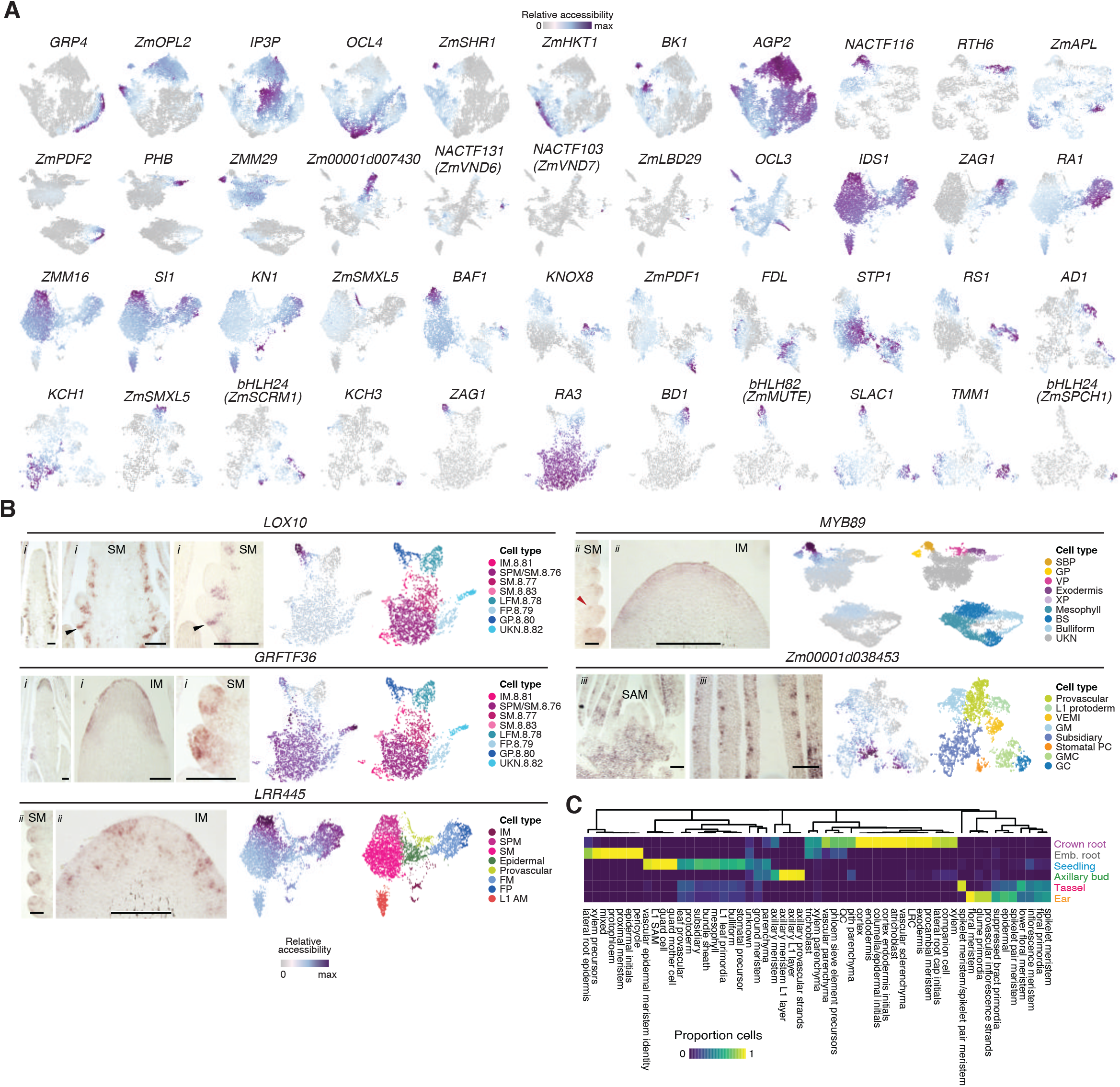
Cell-type annotation and GO enrichment. (A) UMAP embeddings of nuclei barcodes colored by low (grey) to high (dark purple) gene activity values of cell type-specific marker genes. (B) RNA *in situ* hybridization showing expression of *LOX10* in glume primordia and *GRFTF36* in IM and SMs of staminate inflorescence; *LRR445* in the IM periphery and SPMs and *MYB89* in the IM and suppressed bract primordia of pistillate inflorescence; Zm00001d038453 in ground tissue of SAM and leaf primordia sections. Gene accessibility scores and predicted cell types are shown on the right. *i*, tassel primordia. *ii,* ear primordia. *iii*, SAM/leaf. Black triangles point to the glume primordia. Red triangles point to suppressed bract primordia. Size bars illustrate 100-um. AM, axillary meristem; BS, bundle sheath; GC, guard cell; GM, ground meristem; GMC, guard mother cell; GP, glume primordia; IM, inflorescence meristem; L1, layer 1; LFM, lower floral meristem; SAM, shoot apical meristem; SBP, suppressed bract primordia; SM, spikelet meristem; SPM; spikelet pair meristem; Stomatal PC, stomatal precursor; UKN, unknown; VEMI, vascular/epidermal meristematic identity; VP, vascular parenchyma; XP, xylem parenchyma. (C) Proportion of cells within subcluster (column) derived from one of six organs (rows).

**Figure S6.**
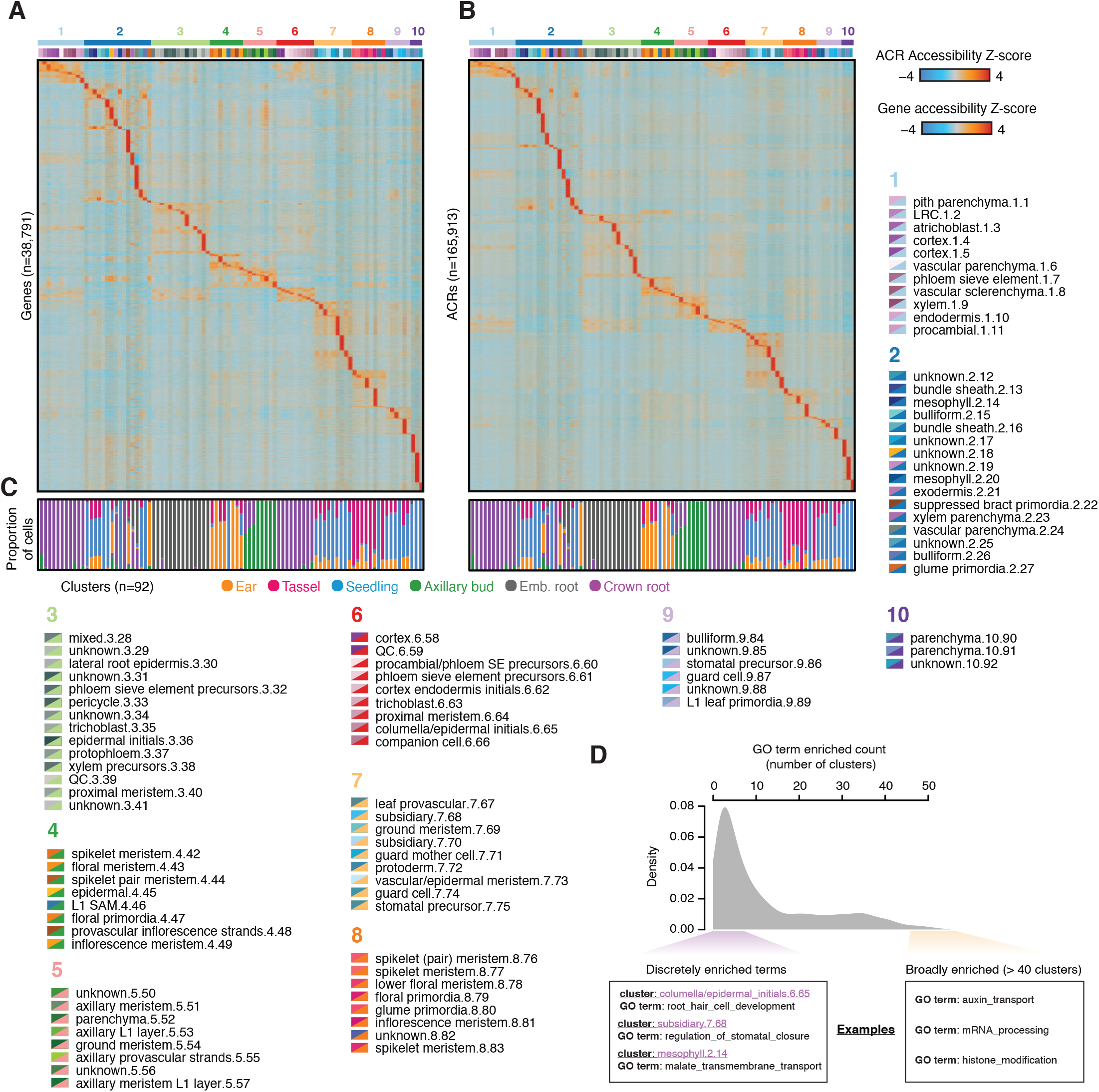
Chromatin accessibility variation across plant cell types. (A) Row gene accessibility Z-scores across cell types. Cell type and subcluster labels are located adjacent to the heatmap in identical order as the matrix. The first number after the cell-type label indicates the major cluster number, while the second number represents the subcluster ID. Legend colors: top left triangle, subcluster (corresponds to the sub-cluster annotation on heatmap). bottom right triangle, major cluster (corresponds to the major cluster annotation on heatmap). (B) Row ACR chromatin accessibility Z-scores across cell types. (C) Proportion of cells derived from one of six organs within each cell type/subcluster. (D) Distribution of GO term enrichment across clusters, where the x-axis indicates the number of clusters in which a GO term is significantly enriched.

**Figure S7.**
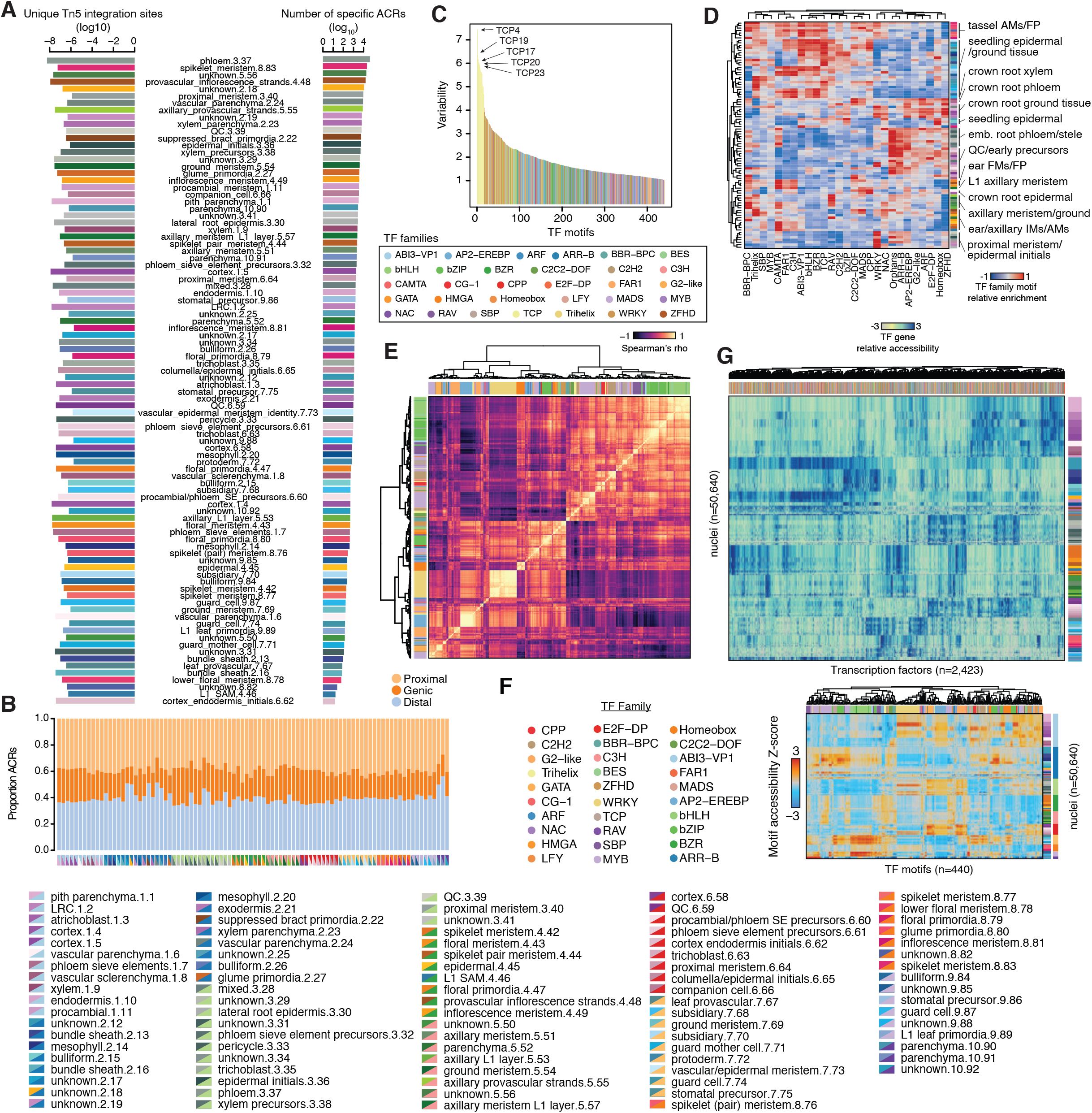
Transcription factor motif variation underly dynamic accessible regions. (A) Comparison of cell type-specific ACR counts (by cell-type) and the number of unique Tn5 integration sites. (B) Proportion of distal, genic and proximal ACRs per cell type. (C) Ranked TF motif variability across cells, colored by TF family. (D) Mean TF family motif enrichment (average deviation scores per cell type per TF family) across cell-types scaled to +/- 1. (E) Spearman’s correlation coefficient (rho) between TF motifs (comparison of motif deviations across all nuclei). Row and column colors represent TF motif families. (F) TF motif deviations for 440 TF motifs (columns) per nucleus (rows). The TF family for each corresponding motif is denoted by column header colors. Cluster identification (sub-cluster/major cluster) are illustrated as row header colors on the right side of the heatmap. (G) Relative gene accessibility scores for 2,423 maize transcription factors (columns) for 50,640 nuclei (rows). Nuclei are sorted by sub-cluster cell type (row color labels).

**Figure S8:**
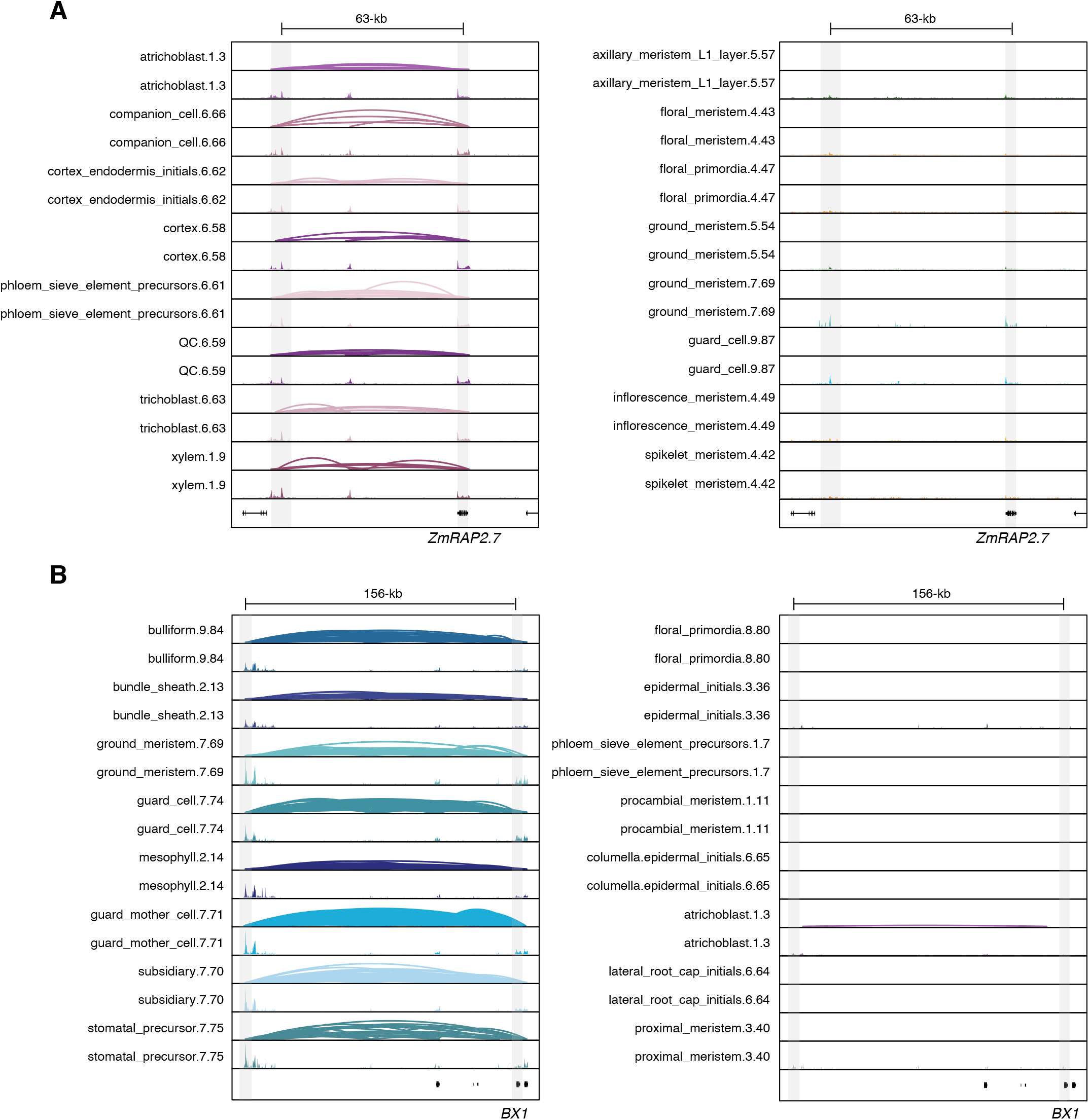
Capture of known long-range chromatin loops in maize. (A) Co-accessible ACRs at the *ZmRAP.2* locus in maize, with root-specific expression patterns, across eight root-derived (left) and eight above-ground (right) cell types. The height of the loops reflects the strength of co-accessibility. Pseudobulk chromatin accessibility tracks are shown under co-accessible ACR linkages for each cell-type. (B) Co-accessible ACRs at the *BX1* locus in maize, predominantly expressed in seedling tissue, across eight seedling-derived (left) and eight non-seedling derived cell types. The height of the loops reflects the strength of co-accessibility. Pseudobulk chromatin accessibility tracks are shown under co-accessible ACR linkages for each cell-type.

**Figure S9.**
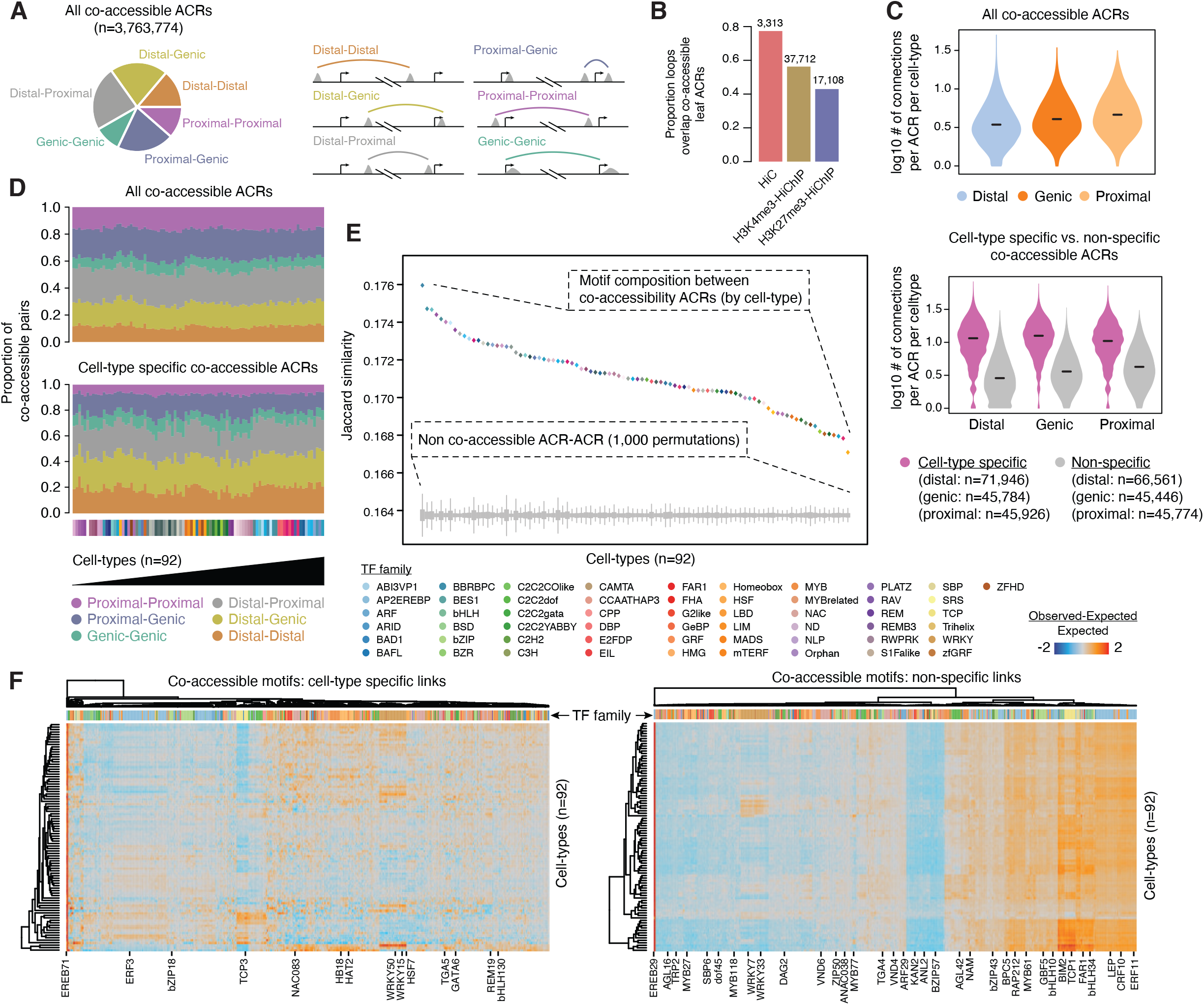
Co-accessible ACRs reflect in vivo chromatin interactions driven by coordinated TF activity. (A) Proportions of co-accessible ACR types, illustrated by toy examples. (B) Proportion leaf Hi-C, H3K4me3-HiChIP and H3K27me3-HiChIP chromatin loops that overlap co-accessible ACRs from leaf cell types (clusters with greater than 50% of cell derived from seedlings). (C) Top, log10 number of connections per ACR per cell type from all co-accessible ACRs split by genomic context: distal, proximal, and genic. Bottom, log_10_ number of connections per ACR per cell type from cell type-specific (purple) and non-specific (grey) co-accessible ACRs, split by genomic context. (D) Proportion of co-accessible classifications by cell type for all co-accessible ACRs (top) and cell type-specific co-accessible ACRs (bottom). (E) Jaccard similarity of motif composition between co-accessible ACR edges by cell type (colored diamonds) relative to the same number of random ACR-ACR links, permuted 1,000 times for each cell-type (grey boxplots). Box plots represent the interquartile range, grey lines indicate the permuted range. (F) Heatmaps of observed proportion of co-accessible ACRs with the same motif embedded within link edges subtracted and divided by the expected proportion estimated by 1,000 permutations using random ACR-ACR links with the same number of co-accessible ACRs.

**Figure S10.**
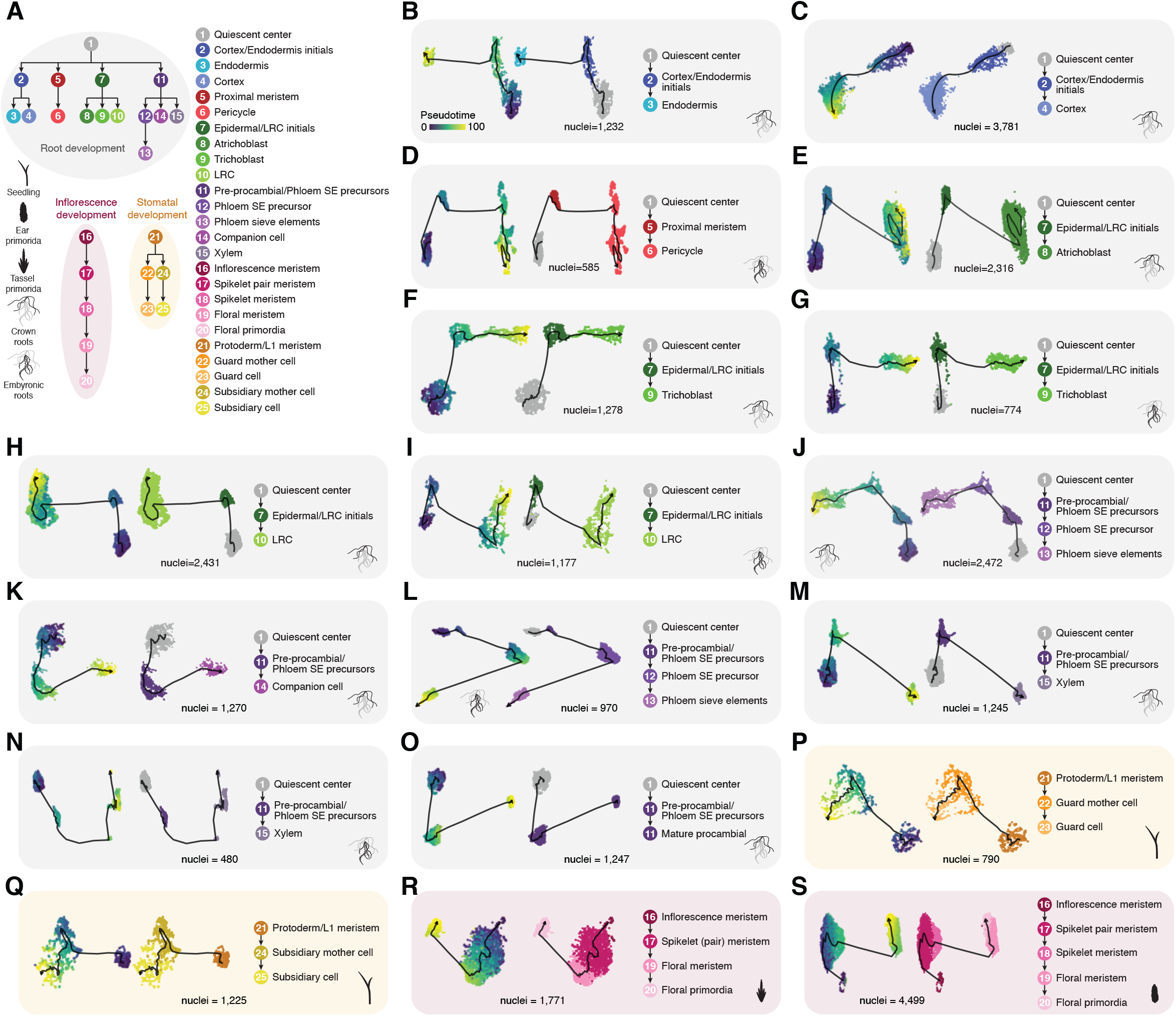
Pseudotime trajectory construction. (A) Overview of pseudotime developmental trajectory analysis from four organs: root, seedling, tassel (staminate inflorescence), and ear (pistilate inflorescence). (B) Endodermis development in crown roots. (C) Cortex development in crown roots. (D) Pericycle development in embryonic roots. (E) Atrichoblast development in crown roots. (F) Trichoblast development in crown roots. (G) Trichoblast development in embryonic roots. (H) Lateral root cap (LRC) development in crown roots. (I) Lateral root cap (LRC) development in embryonic roots. (J) Phloem sieve element (SE) development in crown roots. (K) Companion cell development in crown roots. (L) Phloem sieve element (SE) development in embryonic roots. (M) Xylem development in crown roots. (N) Xylem development in embryonic roots. (O) Procambial development in crown roots. (P) Guard cell development in seedling. (Q) Subsidiary cell development in seedlings. (R) Floral primordia development in staminate inflorescence (tassel). (S) Floral primordia development in pistillate inflorescence (ear).

**Figure S11.**
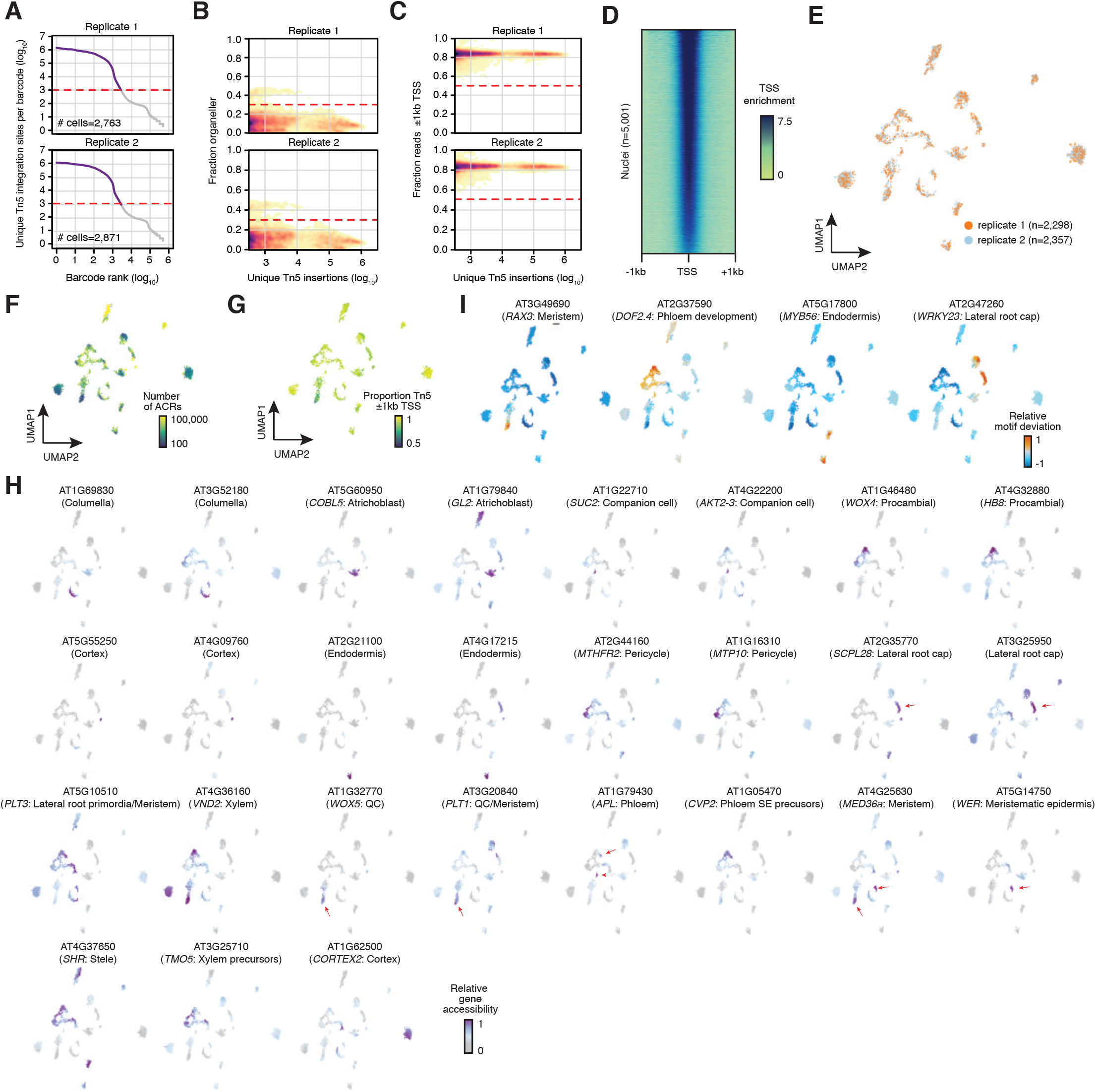
*Arabidopsis thaliana* root cell type atlas. (A) Knee plots for Arabidopsis thaliana root samples illustrating log_10_ transformed cellular read depths of log_10_ ranked barcodes across two biological replicates. (B) Density scatter plots of log_10_ transformed barcode read depths (x-axis) by the fraction of Tn5 integration sites derived from organellar sequences (chloroplast and mitochondrial) relative to the total number of unique Tn5 integration sites associated with each barcode from the two biological replicates. Dashed red lines indicate the threshold of two standard deviations from the mean used to filter lower quality barcodes. (C) Density scatter plots of log_10_ transformed barcode read depths (x-axis) by the fraction of Tn5 integration sites mapping to within 2-kb of transcription start sites (TSSs). Dashed red lines indicate the threshold of two standard deviations from the mean used to filter lower quality barcodes. (D) Average TSS enrichment (normalized read depth adjusted by the two 10 bp windows 1-kb away from TSSs) across 5,001 *Arabidopsis thaliana* root barcodes (rows). (E) UMAP (Uniform manifold approximation projection) embeddings of *Arabidopsis thaliana* root barcodes colored by biological replicate. (F) total number of accessible chromatin regions (ACRs); (G) the proportion of Tn5 integration sites within 1-kb of TSSs. (H) Relative gene accessibility for 27 known cell-type/domain restricted marker genes used to inform cell-type annotation of Arabidopsis thaliana root clusters. (I) Relative motif deviations for transcription factors with known cell-type specificities.

**Figure S12.**
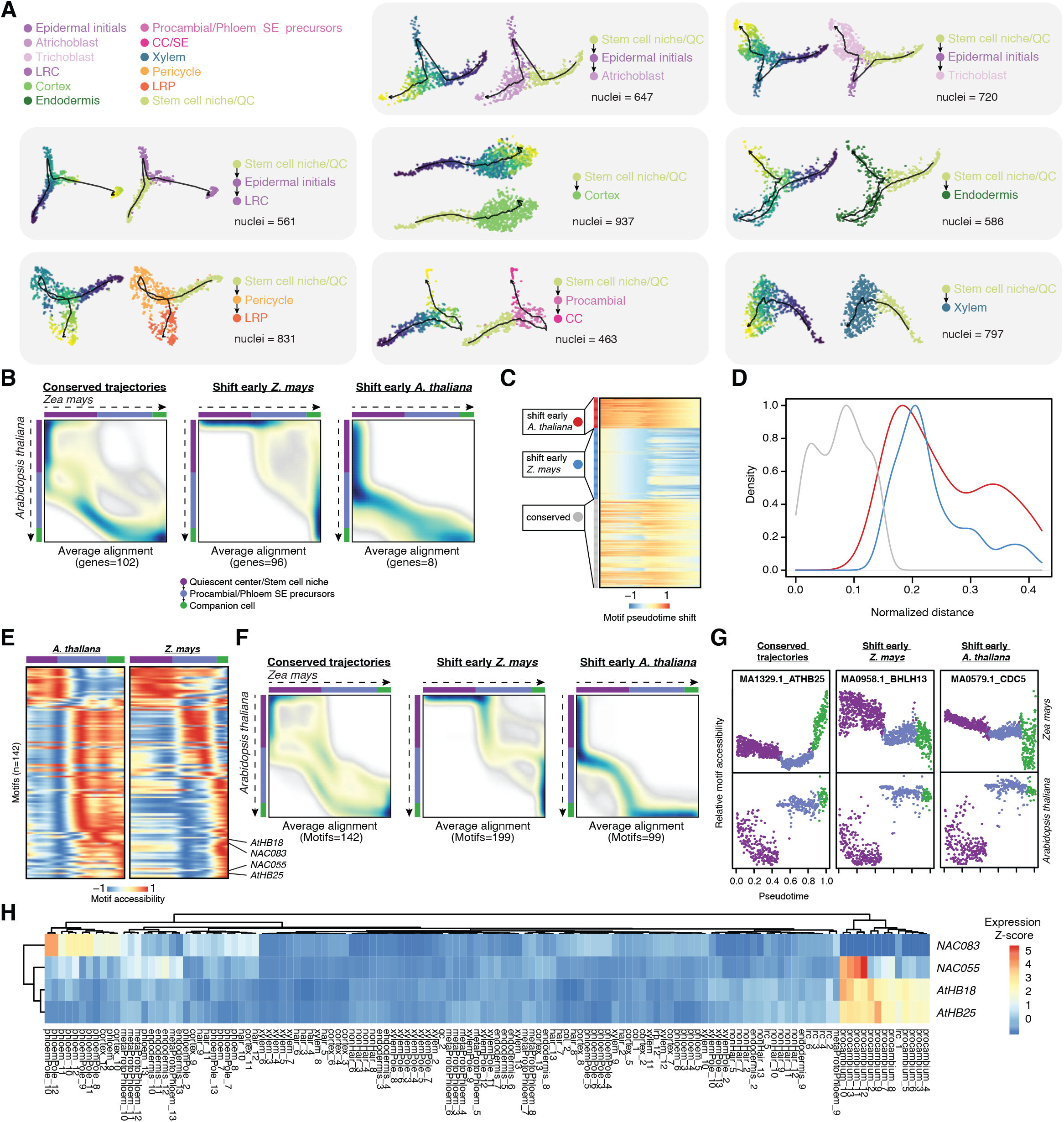
Dynamic and conserved chromatin accessibility across pseudotime between Arabidopsis thaliana *and* Zea mays. (A) Pseudotime trajectories for Atrichoblast, Trichoblast, Lateral root cap (LRC), Cortex, Endodermis, Lateral root primordia (LRP), Companion cells (CC), and Xylem development. (B) Averaged alignments of conserved, shift early *Z. mays*, and shift early *A. thaliana* putative orthologs. (C) Pseudotime shifts of TF motifs between *A. thaliana* and *Z. mays*, clustered into k-means and conserved groups. (D) Distributions of motif-motif normalized distances between *Z. mays* and *A. thaliana* for the three groups. (E) Conserved motifs (n=142) ordered by pseudotime. Heatmaps for *A. thaliana* and *Z. mays* have identical row orders. (F) Averaged alignments of conserved, shift early *Z. mays*, and shift early *A. thaliana* groups based on motif-motif global alignments from the dynamic time-warping algorithm. (G) Examples of conserved, shift early *Z. mays*, and shift early *A. thaliana* motifs from both species. (H) Gene expression Z-scores across *A. thaliana* FAC sorted root cell-types for the TFs recognizing the top four conserved motifs.

## STAR*METHODS

### RESOURCE AVAILABILITY

#### Lead Contact

Further information and requests for resources and reagents should be directed to and will be fulfilled by the Lead Contact, Bob Schmitz.

#### Materials Availability

This study did not generate new unique reagents.

#### Data and Code Availability

Raw and processed data has been deposited in NCBI GEO database under accession code GSE155178. Code used throughout the analysis can be found in the following GitHub repository: https://github.com/plantformatics/maize_single_cell_cis_regulatory_atlas. We also released an R package for pre-processing, normalization, clustering, and other downstream analytical steps into streamlined toolkit of scATAC-seq data can be found in the following GitHub repository: https://github.com/plantformatics/Socrates.

### EXPERIMENTAL MODEL AND SUBJECT DETAILS

#### Growth conditions

For libraries derived from seedlings, kernels from genotypes B73 and Mo17 were obtained from USDA National Plant Germplasm System (https://npgsweb.ars-grin.gov) and sown in Sungro Horticulture professional growing mix (Sungro Horticulture Canada Ltd.). Soil was saturated with tap water and placed under a 50/50 mixture of 4100K (Sylvania Supersaver Cool White Delux F34CWX/SS, 34W) and 3000K (GE Ecolux w/ starcoat, F40CX30ECO, 40W) lighting. Seedlings were grown under a photoperiod of 16 hours of light, eight hours of dark. The temperature was approximately 25°C during light hours with a relative humidity of approximately 54%.

#### Maize seedlings

Above ground seedling tissues were harvested between 8 and 9 AM six days (V1-stage) after sowing. We used both fresh (B73/Mo17 pooled) and flash frozen (B73 only) seedling tissue to construct scATAC-seq libraries (**Table S1**).

#### Maize roots

Maize root samples were obtained as follows: B73 kernels were sterilized with 70% EtOH treatment for 5 minutes. After removing the ethanol solution, kernels were suspended with 50% bleach for 30 minutes, followed by five washes with autoclaved Milli-Q water. Sterilized kernels were then sown onto mesh plates with half strength MS (Phytotech laboratories, catalog: M519) media and wrapped in Millipore tape. Plates were incubated in a Percival growth chamber with a photoperiod of 16 hours of light, eight hours of dark. The growth chamber temperature was set to 25°C with a relative humidity of approximately 60%. Apical root tips (bottom 2 cm) of seminal and primary root samples were harvested six days (V1-stage) after sowing between 8 and 9 am. Crown root samples (21 days after sowing) were derived from the three developmental zones of greenhouse grown B73 plants between 8 and 9 am and rinsed with sterile water 3 times.

#### Maize Inflorescence

Data generated from young inflorescence (ear and tassel primordia) were derived from B73 maize grown in the greenhouse. Inflorescence primordia were extracted from shoots harvested approximately one month (V7-stage, 2-4 mm) after sowing, between 8 and 9 AM. Inflorescence primordia between three and eight millimeters from the base to the apical tip were placed in sterile water and used for nuclei isolation.

#### Maize axillary buds

Axillary buds (∼30 samples per library) were taken from B73 maize plants grown in the greenhouse at approximately the same developmental stage (V7) as tassel and ear primordia.

#### Arabidopsis roots

Seven-day old *A. thaliana* roots were prepared similarly as for maize with the exception of deriving nuclei from whole roots.

### METHOD DETAILS

#### Single cell ATAC-seq library preparation

Each library was prepared by mixing at least three independent biological samples (3-4 seedlings, 3 tassel or ear primordia, 12-14 root tips, 12-14 crown root samples, ∼30 axillary buds, and 100-200 *A. thaliana* whole roots). One scATAC-seq library (B73 seedling) was derived from flash frozen tissue (liquid nitrogen, followed by 7-day −80°C storage), while the remaining libraries were constructed with freshly harvested tissue (**Table S1**).

To isolate individual plant nuclei, fresh or flash frozen tissue from multiple biological samples were placed on petri dishes and vigorous chopped with a No. 2 razor blade for two minutes in ∼500 uL LB01 buffer (15mM Tris pH 7.5, 2mM EDTA, 0.5mM Spermine, 80mM KCl, 20mM NaCl, 15mM 2-ME, 0.15% TrixtonX-100). Homogenized tissue was then filtered through two layers of miracloth, stained with DAPI to a final concentration of ∼1uM and loaded onto a Beckman Coulter MoFlo XDP flow cytometer instrument. A total of 120,000 nuclei were sorted for each sample across four catch tubes (30,000 nuclei each) containing 200 uL LB01. Isolated nuclei were spun down in a swinging-bucket (5 minutes, 500 rcf) centrifuge resuspended in 10uL LB01, pooled, and then visualized on a hemocytometer with a fluorescent microscope. Nuclei suspensions were then spun down (5 minutes, 500 rcf) and resuspended in diluted nuclei buffer (10X Genomics) to a final concentration of 3,200 nuclei per uL and used as input for scATAC-seq library preparation (5 uL; 16,000 nuclei total). Samples were kept on ice for all intermittent steps. For B73/Mo17 mixed library, we pooled 8,000 nuclei from both B73 and Mo17 that were independently isolated. Single-cell ATAC-seq libraries were constructed according to the manufacture’s instruction (10X Genomics, catalog: 1000176). Libraries were sequenced with Illumina NovaSeq 6000 in dual-index mode with eight and 16 cycles for i7 and i5 index, respectively.

#### Single nuclei RNA-seq library preparation

We prepared snRNA-seq libraries from two biological replicates, each composed of three independent 7-day old B73 seedlings. Seedlings were vigorously chopped with a No. 2 razor blade on a petri dish in 500 uL of nuclei isolation buffer (Phosphate-Buffered Saline [PBS; ThermoFisher], 500U SUPERase RNase inhibitor [Invitrogen], 1mM 1,4-Dithiothreitol [DTT; Millipore Sigma], and 0.05% Triton X-100 [Millipore Sigma]). Homogenized tissue in nuclei isolation buffer was filtered through a 40-um cell strainer (pluriSelect) and spun at 500 rcf for 5 minutes. The supernatant was discarded, followed by two more wash (500 uL nuclei isolation buffer) and centrifugation steps (500 rcf for 5 minutes), discarding the supernatant and resuspending in 10 uL nuclei isolation buffer lacking Triton X-100. The concentration of nuclei in solution was estimated on a hemocytometer under a fluorescent microscope and adjusted to 2,000 nuclei per uL with nuclease-free water. Single-nuclei RNA-seq libraries were prepared from a total of 16,000 nuclei per library following the manufactures instructions for the Single Cell Gene Expression 3’ V3 library kit (10X Genomics, catalog: 1000269). Libraries were sequenced on an Illumina NovaSeq 6000 in dual-index mode.

#### In situ hybridizations

3-4mm tassel and ear primordia and young seedlings from the maize B73 inbred line were dissected and fixed in a cold paraformaldehyde acetic acid solution (4% PFA) for 48 hours. Following dehydration through a graded ethanol series and clearing of the tissue with a Histo-clear II solution (Electron Microscopy Sciences), samples were embedded using Paraplast Plus tissue embedding media (McCormick Scientific). 8mm sections were hybridized at 56°C with antisense probes labelled with digoxigenin (DIG RNA labeling mix, Roche), and detected using NBT/BCIP (Roche). Probes were synthesized by *in vitro* transcription (T7 RNA polymerase, Promega) of PCR products obtained from embryo cDNA or from digested full-length cDNA clones. The vectors and primers used for probe design are listed in **Table S18**.

### QUANTIFICATION AND STATISTICAL ANALYSIS

#### scATAC-seq raw reads processing

The following data processing was performed using each tissue and/or replicate independently unless noted otherwise. Raw BCL files were demultiplexed and convert into fastq format using the default settings of the 10X Genomics tool *cellranger-atac makefastq* (v1.2.0). Partial raw read processing (adapter/quality trimming, mapping and barcode attachment/correction) was carried out with *cellranger-atac count* (v1.2.0) using AGPv4 of the maize B73 reference genome (Jiao et al., 2017). Properly paired, uniquely mapped reads with mapping quality greater than 10 were retained using *samtools view* (v1.6; -f 3 -q 10) and by filtering reads with XA tags (Li et al., 2009). Duplicate fragments were collapsed on a per-nucleus basis using *picardtools* (http://broadinstitute.github.io/picard) *MarkDuplicates* (v2.16; BARCODE_TAG=CB REMOVE_DUPLICATES=TRUE). Reads mapping to mitochondrial and chloroplast genomes were counted for each barcode, then excluded from downstream analysis. We removed reads representing potential artifacts by excluding alignments coincident with a blacklist of regions composed of low-complexity and homopolymeric sequences (*RepeatMasker* v4.07) (AFA Smit, 2013-2015), nuclear sequences with homology (greater than 80% identity and coverage) to mitochondrial and chloroplast genomes (*BLAST+* v2.7.1) (Camacho et al., 2009), regions exhibiting Tn5 integration bias from Tn5-treated genomic DNA (1-kb windows with greater than 2-fold coverage over the genome-wide median), and potential collapsed sequences in the reference (1-kb windows with greater than 2-fold coverage over the genome-wide median using ChIP-seq input). Genomic Tn5 and ChIP input data were acquired from Ricci, Lu and Ji et al. BAM alignments were then converted to single base-pair Tn5 integration sites in BED format by adjusting coordinates of reads mapping to positive and negative strands by +4 and −5, respectively, and retaining only unique Tn5 integration sites for each distinct barcode. Sequencing saturation was calculated as the proportion of unique reads relative to the estimated library complexity output by the *MarkDuplicates* function apart of picardtools.

#### Cell calling

To identify high-quality nuclei (a term used interchangeably with “barcodes”) using the filtered set of alignments, we implemented heuristic cutoffs for genomic context and sequencing depth indicative of high-quality nuclei. Specifically, we fit a smoothed spline to the log_10_ transformed unique Tn5 integration sites per nucleus (response) against the ordered log_10_ barcode rank (decreasing per-nucleus unique Tn5 integration site counts) using the *smooth.spline* function (spar=0.01) from base R (Team, 2013). We then used the fitted values from the smoothed spline model to estimate the first derivative (slope), taking the local minima within the first 16,000 barcodes as a potential knee/inflection point (16,000 was selected to match the maximum number of input nuclei). We set the unique Tn5 library depth threshold to the lesser of 1,000 reads and the knee/inflection point, excluding all barcodes below the threshold. Spurious integration patterns throughout the genome can be representative of incomplete Tn5 integration, fragmented/low-quality nuclei, or poor sequence recovery, among other sources of technical noise. In contrast, high quality nuclei often demonstrate a strong aggregate accessibility signal near TSSs. Therefore, we implemented two approaches for estimating signal-noise ratios in our scATAC-seq data. First, nuclei below two standard deviations from the mean fraction of reads mapping to within 2-kb of TSSs were removed on a per-library basis. Then, we estimated TSS enrichment scores by calculating the average per-bp coverage of 2-kb windows surrounding TSSs, scaling by the average per-bp coverage of the first and last 100-bp in the window (background estimate; average of 1-100-bp and 1901-2000-bp), and smoothing the scaled signal with rolling-means (R package; *Zoo*). Per barcode TSS enrichment scores were taken as the maximum signal within 250-bp of the TSS. Lastly, for each library, we removed any barcode with a proportion of reads mapping to chloroplast and mitochondrial genomes greater than two standard deviations from the mean of the library.

#### Detection of multiplet droplets

To estimate the empirical proportion of doublets present in our data, we demultiplexed the two-genotype (B73 and Mo17) pooled seedling scATAC-seq sample and assessed the proportion of barcodes reflecting a mixtures of reads derived from both genotypes. Specifically, B73 and Mo17 whole genome short read resequencing data were acquired from PRJNA338953. Paired-end reads were quality and adapter trimmed with *fastp* (v0.19.5) (Chen et al., 2018) and aligned to the B73 v4 maize reference genome (Jiao et al., 2017) using *BWA mem* (Li, 2013) with non-default settings (-MT 1). Duplicate reads were removed using *samtools rmdup* (Li et al., 2009) (v1.6). The genomic coordinates of short nucleotide variants (SNVs; single nucleotide polymorphisms [SNPs] and small insertions/deletions [INDELs]) for both genotypes were identified using *freebayes* (Garrison and Marth, 2012) (v1.0.0) with non-default settings (--min-repeat-entropy 1 --min-alternate-fraction 0.05). Only biallelic SNPs – requiring at least 5 reads per genotype where B73 and Mo17 were homozygous for reference and alternate nucleotides, respectively – were retained. Genotypes were called by modeling allele counts as a binomial distribution with a term accounting for the sequencing error rate, *E_t_* (determined empirically as the fraction of SNPs failing to match either allele), estimating posterior probabilities via Bayes theorem, and assigning the genotype (or mixture of genotypes) with the greatest probability (Eq. 1–7). Specifically, the probability to observe *k* out of *n* SNPs from B73 can be modeled as a binomial distribution for each B73 (*A_1_*), Mo17 (*A_2_*), and doublet barcode state (*N*) (Eq. 1–3):

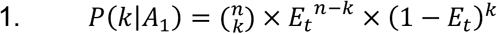

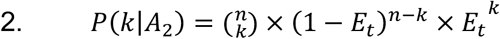

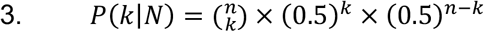

Let *p*(*A*_1_|*k*), *p*(*A*_2_|*k*), and *p*(*N*|*k*) reflect posterior probabilities for genotypes B73, Mo17, and doublet barcodes given *k* allele counts from B73; posterior probabilities can be estimated as follows (Eq. 4–6):

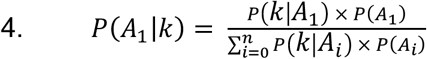

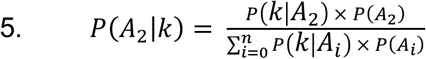

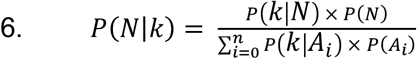

Finally, the genotype called for each barcode was determined as the event with the greatest posterior probability (Eq. 7):

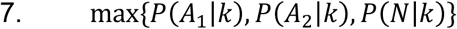

#### In silico sorting

To provide sufficient sensitivity for peak calling prior to clustering, we followed an *in-silico* sorting strategy to identify crude clusters of similar cells within each organ (Cusanovich et al., 2018). To do so, we generate a binary matrix representing the presence/absence of Tn5 integration sites in 1-kb windows across all cells in a given organ. Bins with less than 1% accessible cells and cells with less than 100 accessible bins were removed. This binary matrix was then transformed using the matrix normalization method term-frequency inverse document-frequency (TF-IDF). Briefly, the TF term was estimated by weighting binary counts at each bin by the total number of bins containing Tn5 integration sites in a given cell, scaling each cell to sum to 100,000, adding a pseudo-count of one, and log transforming the resulting values to reduce the effects of outliers in downstream processing. The IDF term was calculated as the log transformed ratio of the total number of nuclei to the number of nuclei that were marked as accessible for a given bin. We add a pseudo-count of one to the inverse frequency term to avoid taking the log of zero. The TF-IDF scaled matrix was estimated by taking the dot product of the TF and IDF matrices. To enable faster downstream computation, we kept the top 25,000 bins with the greatest TF-IDF variance across nuclei. The reduced TF-IDF matrix was denoised with singular value decomposition (SVD), retaining the 2^nd^ – 11^th^ dimensions (termed Latent Semantic Indexing, LSI). Each row was centered and standardized, capping the values at ± 1.5. Crude clusters were visually identified using ward.D2 hierarchical bi-clustering on the cosine distances of LSI nuclei and bin embeddings.

#### ACR identification

ACRs were identified by treating each bulk and single-cell ATAC-seq library as a traditional bulk ATAC-seq library. Aligned reads were filtered by mapping quality greater than 10, and duplicate reads were removed via *samtools rmdup.* We then identified ACRs for each library by converting the BAM alignments in BED format, adjusting the coordinates to reflect single-base Tn5 integrations, and running *MACS2* (Zhang et al., 2008) with non-default parameters: --extsize 150 --shift -75 --nomodel --keep-dup all. A final set of ACRs for comparing bulk and aggregate scATAC-seq libraries (**Figure S1**) was constructed by taking the union of ACRs across all libraries. To leverage the increased sensitivity afforded by cell-type resolved cluster information while ensuring robust reproducibility in ACR identification, we generated pseudo-replicated bulk alignments using the LSI-based crude clusters (see above, “*In-silico* sorting”). Pseudo-replicates were constructed by randomly allocating nuclei from each cluster into two groups, with a third group composed of all cells from the cluster (cluster bulk). These groupings were used to concatenate Tn5 integration sites corresponding to the nuclei from each group into three BED files. ACRs were then identified from the enrichment of Tn5 integration sites from the pseudo-replicate or cluster bulk aggregates using *MACS2* run with non-default parameters: --extsize 150 --shift -75 --nomodel --keep-dup all. ACRs from both pseudo-replicates and the cluster bulk were intersected with BEDtools, retaining ACRs on the conditional intersection of all three groupings (both pseudo-replicates and the cluster bulk) by at least 25% overlap. The remaining ACRs were then redefined as 500-bp windows centered on the ACR coverage summit. To integrate information across all clusters, ACRs from each cluster were concatenated into a single master list. Lastly, overlapping ACRs were filtered recursively to retain the ACR with the greater normalized kernel Tn5 integration density as previously described (Satpathy et al., 2019).

#### Nuclei clustering

Starting with a binary nucleus x ACR matrix, we first removed ACRs that were accessible in less than 0.5% of all nuclei, and filtered nuclei less than 50 accessible ACRs. Inspired by recent developments in modeling single-cell RNA-seq data (Hafemeister and Satija, 2019), we developed a regularized quasibinomial logistic framework that overcomes noise inherent to sparse, binary scATAC-seq data by pooling information across ACRs while simultaneously removing variation due to technical effects, particularly those stemming from differences in barcode sequencing depths. First, a subset of 5,000 representative ACRs selected by kernel density sampling of ACR usage (fraction nuclei that are accessible at a given ACR) were used to model the parameters of each ACR, using ACR usage as a covariate in a generalized linear model. Specifically, the expected accessibility of an ACR, *y_i_*, can be estimated with a generalized linear model containing a binomial error distribution and logit-link function, and an overdispersion term with a quasibinomial probability density function (Eq. 8).

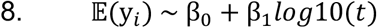

Where *t* is a vector of the sums of accessible ACRs across cell *j* (Eq. 9):

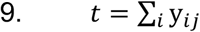

To prevent over-fitting and ensure robust estimates in light of sampling noise, we learned the global regularized model parameters, including overdispersion, using the representative ACRs by fitting each parameter against the log_10_ fraction of accessible nuclei via kernel regression, resulting in smoothed parameter estimates across the spectrum of ACR accessibility penetrance present in these data. The learned global regularized model parameters were then used to constrain fitted values across all ACRs for each nucleus with a simple affine transformation. To account for technical variation among nuclei (variation in barcode log_10_ transformed read-depth, in particular) we calculated Pearson residuals for each ACR, scaling the residuals by the regularized dispersion estimate and centering values via mean subtraction, representing variance-stabilized and read-depth normalized values of accessibility for a nucleus at a given ACR. We note that this method is amenable to calculating residuals that account for additional sources of technical variation, including categorical and numeric covariates, that may obscure biological signal, such as batch effects, proportion of mitochondrial reads, etc.

The dimensionality of the Pearson residual matrix was reduced using singular value decomposition (SVD) implemented by the R package *irlba* (Witten et al., 2009), retaining the first 25 left singular vectors scaled by singular values (hereafter referred to as nuclei embeddings), analogous to principal components (PCs) on an uncentered matrix. Nuclei embeddings were then standardized across components and filtered to remove components correlated with barcode read depth (Spearman’s rho > 0.7). We further reduced the dimensionality of the nuclei embedding with Uniform Manifold Approximation Projection (UMAP) via the R implementation of *umap-learn* (min_dist = 0.1, k=50, metric=”euclidean”). Nuclei were clustered with the *Seurat* v3 (Stuart et al., 2019) framework and Louvain clustering on a k=50 nearest neighborhood graph at a resolution of 0.02 with 100 iterations and 100 random starts. Clusters with aggregated read depths less than 1.5M were removed. To filter outliers in the UMAP embedding, we estimated the mean distance for each nucleus with its *k* (*k*=50) nearest neighbors and removed nuclei greater than 3 standard deviations from the mean.

We observed fine-scale heterogeneity within major clusters, thus we repeated our clustering pipeline for each major cluster independently by partitioning the SVD embedding into the top 20 components, L2 normalizing nuclei embeddings across components, and projecting the L2-normalized embeddings into the UMAP space. Subclusters of nuclei were identified by Louvain clustering on the L2 normalized SVD embedding (resolution set manually, range = 0.6 – 1.0) with 20 nearest neighbors, filtering outlier nuclei more than 2 standard deviations from the mean distance of 25 nearest neighbors within each cluster.

For analysis of chromatin accessibility across clusters, we assembled a matrix of clusters by ACRs by aggregating the number of single-base resolution Tn5 integration sites from nuclei within the same cluster for each ACR, analogous to normalizing by the proportion of reads in peaks for each cluster. To account for differences in read depth and other technical factors, the raw counts were transformed with *edgeR’s* “cpm” (log=T, prior.count=5) as previously described (Corces et al., 2018). Log-transformed ACR coverage scores were quantile normalized using “normalize.quantiles” with the R package, *preprocessCore.* Finally, to aid data visualization, we estimated per ACR Z-scores across clusters by mean subtraction and standardization (identical to row-wise execution of the R function, “scale”).

#### Identification of co-accessible ACRs

Recent experiments of population-level chromatin accessibility found that pairwise correlations of accessibility among ACRs recapitulates higher-order chromatin interactions observed in Hi-C and other chromatin architecture experiments (Gate et al., 2018). A similar framework was applied to populations of single cells, which showed that co-accessible ACRs are typically more conserved and functionally associated (Buenrostro et al., 2015). To identify potentially functional co-accessible ACRs, we applied a recently developed method, *Cicero* (Pliner et al., 2018), that estimates regularized correlation scores (ranging from −1 to 1) among nearby ACRs with graphical LASSO to penalize potential interactions by physical distances. Using the binary nuclei x ACR matrix as input, we subset nuclei by their subcluster IDs and estimated co-accessibility among ACRs within 500-kb for each of the 92 clusters, independently. *Cicero* was run by applying a background sample of 100 random regions, and 15 nuclei pseudo-aggregates based on k-nearest-neighbors derived from the UMAP coordinates. To control the false discovery rate (FDR) of co-accessible ACR calls, we shuffled the nuclei x ACR matrix such that the total number of reads per ACR and reads per nucleus were identical to the original matrix. We then repeated co-accessible ACR identification with the shuffled matrix, keeping the original parameters to *Cicero* unchanged. Empirical FDR cluster-specific cut-offs were constructed by identifying the minimum positive co-accessibility score in the background where the FDR < 0.05. Co-accessible links below cluster-specific thresholds were removed. Co-accessible ACRs passing thresholds were compared with previously published HiC and HiChIP data sets derived from maize seedling and pistillate inflorescence primordia (Ricci et al., 2019).

#### Estimation of gene accessibility scores

Chromatin accessibility at TSSs and gene bodies exhibit marked correlation with transcription output in bulk samples (**Extended Data Fig. 3f**). To aid the identification of marker genes underlying distinct cell-types, we used *Cicero* to estimate gene activity scores. *Cicero* models gene activity as a weighted accessibility score that integrates both proximal and distal regulatory elements linked to a single gene by co-accessibility analysis (see above section “Identification of co-accessible ACRs”). Relative gene accessibility scores per nucleus were estimated by taking a weighted average (3:1, gene body score to proximal/distal activity) of the scaled number of reads mapping to gene bodies for each barcode (summing to 1) with the *Cicero* estimate of gene activity derived from ACRs mapping to 1-kb upstream of gene TSSs and their associated distal ACRs linked by co-accessible ACRs passing FDR < 0.05 thresholds (connected ACRs were constrained to a minimum and maximum intervening distance of 1- and 500-kb, respectively). These weighted gene accessibility scores were rescaled such that gene accessibility scores for a given nucleus summed to 1.

Relative gene accessibility scores exhibited a bimodal distribution with relative gene accessibility values near zero resembling low or non-expressed genes. We applied a gaussian mixture-model (two distributions) based scaling step per cluster to reduce noise introduced by genes with low gene accessibility. Briefly, the average gene accessibility across nuclei was fit to a two distribution gaussian mixture model in each cluster using the R package *mclust*. We estimated cluster-specific scaling parameters determined as the 5% quantile of non-zero gene accessibility values of genes from the gaussian distribution with the larger mean, for each cluster. This parameter was then used to scale gene accessibility scores for all genes in each nucleus within the cluster. Scaled gene accessibility scores were rounded to the nearest integer and normalized across all nuclei and clusters using nucleus-specific size factors estimated as the total gene accessibility of a nucleus divided by the exponential of the mean of log-transformed gene accessibility sums across nuclei. To aid visualization, we smoothed normalized gene accessibility scores by estimating a diffusion nearest neighbor graph (k=15) using the SVD embedding with 3 steps similar to the *run_MAGIC* function in R package *snapATAC* (Fang et al., 2020). Downstream analyzes based on binarized gene accessibility were conducted by simply converting normalized (non-smoothed) accessibility scores to 1 for all positive values.

#### Cell-type annotation

To identify and annotate cell types for each barcode, we identified marker genes known to localize to discrete cell types or domains expected in the sampled tissues/organs based on extensive review of the literature (**Table S2**). To enable gene accessibility comparisons among clusters, we generated three pseudo-replicates for each cluster by resampling nuclei within the cluster such that all cluster pseudo-replicates contained the mean number of nuclei across clusters (number of nuclei per pseudo-replicate = 552) without replacement when possible. To identify genes with increased accessibility relative to other clusters, we constructed a reference panel with three pseudo-replicates by uniformly sampling nuclei without replacement from each organ (number of nuclei per organ = 92), with a total of 552 nuclei per reference panel pseudo-replicate. Read counts per gene were summed across nuclei within each pseudo-replicate. Using the *DESeq2* R package, we identified genes with significantly different (FDR < 0.01) accessibility profiles between each cluster and the reference panel.

The list of significantly differentially accessible genes was filtered to retain the genes on our list of cell type specific markers. We initially ranked the top three marker genes in each cluster by their test statistics. To account for clusters containing small proportions of contaminating nuclei of a different cell type, we adjusted the test statistics using a previously described method (Cusanovich et al., 2018), effectively scaling marker activity scores by the proportion of nuclei in the cluster that were derived from an organ in which the marker gene E is an expected cell type. Clusters where the top three markers corresponded to the same cell type were annotated with the consensus cell type.

As an independent method for cell-type annotation, we devised a resampling and normalization procedure on the log_2_ fold-change values of marker genes to evaluate cell-type enrichment across all possible cell types for each cluster, normalizing enrichment scores by random permutations accounting for different numbers of markers associated with each cell type. Briefly, starting with differential gene accessibility information for each cluster, we iterated over all cell types, extracting markers associated with the cell type of interest. Then, we summed the log_2_ fold-changes values of all markers and multiplied the sum by the proportion of markers passing heuristic thresholds (fold-change > 2 and FDR < 0.01). This score was subtracted by the average of 1,000 random permuted scores from combinations of markers from the remaining cell types (selecting the same number of random genes as the cell type of interest) and divided by the standard deviation of the permuted scores. Cell-type enrichment scores in each cluster were scaled from zero to one by dividing each cell-type enrichment score by the maximum scores across possible cell types. This approach is effective in normalizing differences arising from varying numbers of markers specified for each cell type. Additionally, cell-type annotation scores for clusters with mixed or unknown identity are approximately equally distributed, thus controlling ascertainment bias stemming from marker gene selection. Stated differently, an advantage of this approach is that clusters corresponding to cell types with few or no markers in the tested list are left unassigned as their enrichment scores do not deviate significantly from background levels. Finally, scaled cell-type enrichment scores greater than 0.9 were taken as possible annotations and intersected with putative cell-type labels from the marker ranking approach described above.

For clusters with ambiguous marker gene labels, we developed a logistic regression classifier to identify putative cell types based on whole-genome gene accessibility scores of well-annotated cells. First, we counted the number of Tn5 integration sites per cell overlapping 2-kb upstream to 500-bp downstream of each gene. Read counts were transformed by trimmed mean of M-values (TMM) to enable intra and inter-nucleus comparisons using *edgeR* (Robinson et al., 2010), scaling gene accessibility scores in each nucleus with counts per million. Next, we estimated cell-type enrichment scores for each nucleus by calculating the mean accessibility scores of markers for a given cell type, subtracting the mean background signal defined as 1,000 sets of averaged randomly sampled genes (each set had the same number of genes as the number of markers), divided by the standard deviation of the background signal. Enrichment scores for each nucleus were transformed into a probability distribution by dividing by the sum of cell-type enrichment scores. For each nucleus, we compared the top two most likely cell types, retaining nuclei where the top predicted cell type had a two-fold greater probability than the next most likely assignment. We used these high-confidence cells to train a regularized logistic multinominal classifier with the R package, *glmnet*. Cell-type classifications with less than 10 nuclei in the training set were excluded. We used a LASSO L1 penalty to regularize the logistic classifier, modeling the training set of nuclei as observations and TMM gene accessibility scores as variables. We balanced observations by weighting by the inverse frequency of cell types in the training set. The model was trained with 10-folds and evaluated by testing a 20% hold-out set of nuclei. The predicted cell type for each nucleus in the atlas was taken as the cell type with the greatest probability if the probability ratio between the best and next best assignment was greater than five-fold, otherwise labeled as ‘unknown’. Using these per-cell assignments, we defined subclusters as the majority cell type if greater than 50% of nuclei in the cluster were in agreement, labelling clusters with two or more majority cell types as ‘mixed’ and all other clusters as ‘unknown’. All cell-type labels from these three automated approaches were manually reviewed by careful evaluation with UMAP gene accessibility score embeddings and cluster aggregated coverages for all marker genes and refined *ad hoc*.

#### Cell-cycle annotation

Cell cycle annotation was performed similarly as cell-type annotation. Briefly, we acquired cell-cycle marker genes from Nelms et al. 2019, selecting 35 markers at random for each cell stage (Nelms and Walbot, 2019). The rationale behind selecting equivalent numbers of markers per stage was to prevent biasing cell cycle annotations to cycle stages with more markers, while 35 markers was the minimum gene count across all stages (mitosis). For each stage, we subset the nuclei by gene accessibility (TMM) matrix by the cognate stage, and summed accessibility scores for each nucleus. This cell-cycle stage score was then standardized using the mean and standard deviation of 1,000 permutation of 35 random cell-cycle stage genes, excluding the focal stage. Z-scores corresponding to each cell-cycle stage were converted into probabilities using the R function *pnorm*. Per nucleus posterior cell-cycle probabilities were estimated using Bayes theorem with each cell-cycle stage prior probability set to 0.2 (1/5, for five stages: G1, G1/S, S, G2/M, M). The cell-cycle stage with the maximum probability was selected as the most likely cell stage. Nuclei with multiple cell-cycle annotations with equal maximum probability were considered “ambiguous”.

#### snRNA-seq data processing

Raw fastq files from each snRNA-seq seedling library (across two biological replicates) were processed with *cellranger count* v4.0 to align reads to AGPv4 of the maize B73 reference genome (Jiao et al., 2017). BAM files were filtered to remove multiple mapping reads using a mapping quality filter selecting reads with MQ greater than or equal to 30. The number of nuclear, organeller and transcript-derived unique molecular identifiers (UMIs) reads for each barcode were tabulated from the filtered BAM file. Barcodes with less than 1,000 total UMIs and less than 500 genes with at least one UMI were removed. We then estimated the Z-score distributions for the proportion of mitochondrial, chloroplast, nuclear, and transcript derived UMIs across barcodes. Barcodes above 1 standard deviation (Z-score less than 1) from the mean proportion of UMIs derived from mitochondrial and chloroplast genomes were removed. Likewise, barcodes below 1 standard deviation from the mean proportion of UMIs derived from the nuclear genome were removed.

#### Integration of scATAC-seq and snRNA-seq data

To integrate scATAC-seq and snRNA-seq data into a shared embedding, we input gene accessibility scores and gene expression values from all seedling-derived nuclei passing quality filters described above using *liger* with the function *createLiger* (Welch et al., 2019). Each data set was normalized, subset by highly variable genes, and scaled using the functions *normalize, selectGenes,* and *scaleNotCenter,* sequentially with default arguments. An integrated non-negative matrix factorization (iNMF) embedding was constructed from the gene by nuclei scATAC-seq and snRNA-seq matrices using *optimizeALS* with default settings (k=20, lambda=5). The iNMF embedding was quantile normalized with *quantile_norm* and non-default settings (do.center=FALSE). Louvain clusters from the normalized iNMF embedding were identified at a resolution of 0.25 with *louvainCluster.* To visualize the integrated assays, we used *runUMAP* with non-default settings (n_neighbors = 20, min_dist = 0.01). Differentially accessible and expressed genes per cluster were identified using *runWilcoxon* requiring FDR less than 0.05 and a log_2_ fold change greater 0.25 using the integrated embedding (both gene accessibility and expression across all co-embedded nuclei), gene accessibility in isolation (scATAC-seq nuclei only), and gene expression in isolation (snRNA-seq nuclei only). Differentially accessible ACRs from the normalized (with *liger* function *normalize*) sparse ACR by nuclei matrix were identified using identical heuristic thresholds as for gene expression and accessibility.

To impute ACR accessibility in snRNA-seq derived-nuclei and gene expression values in scATAC-seq nuclei, we ran *imputeKNN* from the *liger* package using either the scATAC-seq or snRNA-seq nuclei as reference cells. We then used the imputed gene expression and ACR accessibility matrices, constrained to only differentially accessible ACRs (n=55,939), to identify significantly associated gene-to-peak linkages with the *liger* function *linkGenesAndPeaks* with non-default settings (dist = ‘spearman’, alpha = 0.05). To remove potential false positives, we shuffled the imputed ACR and gene nuclei matrices and repeated gene-to-peak linkage identification using the same arguments. We then estimated FDR empirically over a grid of a 100 possible correlation values in both the negative and positive directions by identifying correlation cut-offs that removed 95% of gene-to-peak linkages from the shuffled matrices. We then filtered the non-shuffled gene-to-peak linkages according to the thresholds identified from the empirical FDR estimates.

#### STARR-seq analysis

Single bp-resolution enhancer activities were available from a previous study (Ricci et al., 2019). Enhancer activity (defined as the log_2_ ratio between RNA and DNA input fragments scaled per million) for each ACR was taken as the maximum over the entire ACR. A control set of regions was generated to match each ACR with the following criteria: (*i*) GC content within 5%, (*ii*) physically constrained to within 50-kb of an ACR, and (*iii*) the same length (500-bp) distribution. The same set of control regions was used throughout the analysis.

#### Analysis of differential chromatin accessibility

Next, we implemented a logistic regression framework based on binarized ACR accessibility scores for assessing the importance of each ACR to cluster membership by estimating the likelihood ratio between logistic models with, and without a term for cluster membership. Specifically, for each cluster, we compared binarized ACR accessibility scores to a reference panel of uniformly sampled nuclei from each organ (111 nuclei from each organ) where the total number of reference nuclei was set to the average number of nuclei per cluster (n=666). We then fit two generalized linear logistic regression models (Eq. 10–11), with and without a term for membership to the cluster of interest.

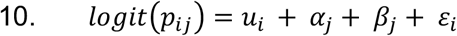

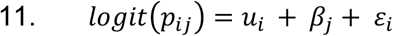

Where *p_ij_* is the probability that ACR *i* is accessible in nucleus *j*, *u_i_* is the proportion of nuclei where ACR *i* is accessible, *α_j_* is the cluster membership of nucleus *j*, *β_j_* is the log_10_ number of accessible ACRs in nucleus *j* and *ε_i_* is the error term for the *i*_th_ ACR. We then used a likelihood ratio test to compare the fits of the two models and estimated the false discovery rate (FDR) using the Benjamini-Hochberg method to identify ACRs that were significantly differentially accessible across clusters by conditioning on FDR < 5% and fold-change threshold greater than two. ACRs meeting these criteria with positive Z-scores in nine or fewer clusters (< 10% of clusters) were considered as cluster-specific. Analysis of differential gene accessibility was performed as described in the section titled “Cell-type annotation”.

#### GO gene set enrichment analysis

Gene set enrichment using GO biological process terms was performed using the R package *fgsea.* For each cluster, test statistics were multiplied by the sign of the log_2_ fold-change value versus the reference panel. GO terms with gene sets less than 10 and greater than 600 were excluded from the analysis. GO terms were considered significantly enriched at FDR < 0.05 following 10,000 permutations.

#### Motif analysis

Motif occurrences were identified genome-wide with *fimo* from the MEME suite toolset (Grant et al., 2011) using position weight matrices (PWM) based on DAP-seq data in *A. thaliana* and *Zea mays* (Galli et al., 2018; O’Malley et al., 2016). To identity TF motifs associated with cell type-specific ACRs, we ranked the top 2,000 ACRs in each cell type by Z-scores derived from CPM normalized accessibility values (see section above “Nuclei clustering”). As a reference for comparison, we identified 2,000 “constitutive” ACRs that varied the least and were broadly accessible across clusters. The number of ACRs containing a specific motif was compared to the frequency of constitutive ACRs harboring the same motif using a binomial test for each cell type and motif. To control for multiple testing, we used the Benjamini-Hochberg method to estimate the FDR, considering tests with FDR < 0.05 as significantly different between the focal cell type and constitutively accessible regions. Maize homologs of *A. thaliana* TFs were identified using protein fasta alignments from BLAST+ v2.10.0 with an E-value cut-off of 1e-5. Only fasta sequences classified as transcription factors from either species were considered during alignment. To narrow the list of putative orthologs based on functional similarity to *A. thaliana* TFs, we filtered alignments with less than 30% identity, removed maize TFs classified as belonging to a different family, and selected the homolog with the greatest Pearson correlation coefficient (PCC) with respect to the motif deviation score. Motif deviation scores of specific TF motifs among nuclei were estimated using *chromVAR* (Schep et al., 2017) with the non-redundant core plant PWM database from JASPAR2018. The input matrix for *chromVar* was filtered to retain a minimum of 50 accessible nuclei per ACR and barcodes with at least 50 accessible ACRs. We visualized differences in global motif usage per nucleus by projecting deviation scores onto the UMAP embeddings. To determine if patterns of TF motif accessibility from individual nuclei could be used to predict cell-type annotations, we constructed a neural network for multinomial classification using the R package, *caret* (Kuhn, 2008) (method=”multinom”) using 80% of nuclei to train, 10-fold cross-validation, averaging error terms across 10 iterations. The nuclei in the 20% withheld group were used to test the model. Sensitivity, specificity and accuracy of the model was evaluated using the function confusionMatrix from *caret*.

To identity *de novo* motifs enriched in accessible but non-transcribed genes, we selected ACRs (n=15,576) within 1-kb of genes that were accessible (ATAC log2 TPM > 1.5) and non-expressed (mRNA log2 TPM < 1) in at least 10 clusters. We then constructed a set of control regions by randomly sampling ACRs within 1-kb of genes expressed (mRNA log2 TPM > 1) and accessible (ATAC log2 TPM > 1.5) in at least 10 clusters (n=15,576). *De novo* motif identification was conducted using the discriminative motif discovery workflow of MEME-ChIP (v5.1.1) with default settings (Machanick and Bailey, 2011). Comparison of *de novo* motifs with experimentally identified motifs was performed using TOMTOM from the MEME-suite toolkit (Gupta et al., 2007).

#### Analysis of cell type-specific selection signatures

Multi-locus allele-frequency differentiation signals between chronologically sampled elite maize inbred lines were mapped onto ACRs (Wang et al., 2020), where the selection score for an ACR was taken as the maximum XP-CLR value within the 500-bp ACR interval. To identify cell types associated with increased signatures of selection, the top 2,000 ACRs defined by standardized quantile-scaled CPM chromatin accessibility (Z-scores, see above “Nuclei clustering”) were identified for each cell type. The mean XP-CLR scores per-cell type were standardized by the mean and standard deviation of randomly sampled ACRs (n=2,000) without replacement across 1,000 permutations, where each permutation estimates the mean XP-CLR scores of a random subset of 2,000 ACRs from the total list of 165,913 possible ACRs. Enrichment Z-scores were converted into *P*-values using the R function *pnorm* (log.p=T, lower.tail=F) and used to estimate FDR via the Benjamini-Hochberg method with the R function *p.adjust* (method=”fdr”).

#### Analysis of co-accessible ACRs

To enable comparison with previously identified Hi-C and HiChIP loops (Ricci et al., 2019), we constrained the distance between co-accessible ACRs to the same range as loops identified in leaf Hi-C and HiChIP (minimum loop distance = 20-kb). Co-accessible ACRs and Hi-C/HiChIP loops were considered overlapping if both anchors overlapped by at least 50-bp. We compared motif composition of co-accessible ACRs by scoring motif occurrence as binary for each ACR and estimating a Jaccard similarity score on the union of motif sets. Motif similarity scores for co-accessible ACRs in each cell type were compared to a null distribution by repeating Jaccard similarity calculations for non-co-accessible ACR-ACR connections (constraining the null connections to blocks of 1,000 ACR on the same chromosome with the same ACR-ACR distance distribution as co-accessible ACRs) across 1,000 permutations. To identify motifs enriched at co-accessible ACR anchors, we first estimated the proportion of co-accessible ACRs with an identical motif at both anchors for each motif and cell type. Then, we constructed the same number of random ACR-ACR connections as co-accessible ACRs, again estimating the proportion of links with an identical motif at both anchors, building a null distribution over 1,000 random permutations. The estimated proportion of co-accessible ACRs with identical motifs at both anchors for each motif was transformed to a Z-score by subtracting and scaling by the mean and standard deviation of the null distribution. Z-scores were converted to *P*-values using the R function, *qnorm* with non-default parameters (log.p=T, lower.tail=F). FDR values were estimated using *p.adjust* (method=”fdr”). Co-accessible motif scores were plotted as heatmaps using *heatmap.2* by subtracting and dividing observed with expected proportions. Rows and columns were clustered with *hclust* (method=”ward.D2”).

#### Pseudotime analysis

Pseudotime trajectories were constructed similar to previous methods (Granja et al., 2020). Briefly, nuclei were ordered based on the principal component space by fitting a continuous trajectory via a smooth spline on the Euclidean distances of each nuclei to a predefined order of cell types. For feature analysis (ACRs, motifs, and TF activity) across pseudotime, nuclei were sorted by ascending pseudotime. The ACR x nucleus matrix was filtered to retain differentially accessible ACRs (see section “Analysis of differential accessibility across pseudotime” below) with at least one nucleus defined as accessible. For each ACR, we fit a generalized additive model with the binary accessibility scores as the response and a smoothed pseudotime component as the dependent variable [s(pseudotime, bs=”cs”)] with a binomial error term and a logit-link function with *gam* from the *mgcv* R package. Predicted accessibility scores across pseudotime were generated from 500 equally spaced interpolated points covering the range of pseudotime values. Finally, predicted accessibility scores were mean-centered, standardized and constrained to the range ±1 for each ACR. Model specification for motif deviations and TF gene accessibility analysis was similar to ACR pseudotime analysis with the exception of a Gaussian error distribution, and TF gene accessibility was normalized by the row maximum rather than rescaling on a ±1 distribution.

#### Analysis of differential accessibility across pseudotime

To identity differentially accessible ACRs across pseudotime, we fit normalized accessibility residuals (Pearson residuals from a generalized linear logistic regression model with log_10_ number of accessible ACRs per barcode as the dependent variable, see section “Nuclei Clustering” above) as the response and pseudotime as the dependent variable using a natural spline with six degrees of freedom [ns(pseudotime, df=6)] from the R package *splines* for each trajectory. We took an F-test based approach for hypothesis testing of differential accessibility across pseudotime by comparing the variance explained by the splined linear model with that of the residuals normalized by degrees of freedom. *P*-values from the model were used to estimate Benjamini-Hochberg FDR values with the R function *p.adjust* (method=”fdr”), where a FDR threshold < 0.05 denoted statistical significance for differentially accessible ACRs across pseudotime. To identify genes and TF motifs with differential accessibility across pseudotime, we fit the linear splined regression model with the normalized gene accessibility scores and motif deviations from each nucleus, respectively, similar to the analysis of ACRs.

#### A. thaliana scATAC-seq processing

scATAC-seq data derived from *A. thaliana* root nuclei were processed similarly to the scATAC-seq data derived from maize nuclei. Specifically, we processed raw fastq files using *cellranger-atac*, filtered multi-mapped reads (MQ less than 10 and the presence XA:Z: tags), removed PCR duplicates by barcode, filtered barcodes by proportion of Tn5 integration sites mapping to organeller genomes above 1 standard deviation from the mean, and removed barcodes with less than 1000 unique Tn5 integration sites. We used *in silico* sorting to group nuclei by similarity, identify ACRs, estimate residuals with regularized quasibinomial regression from the binary ACR by nuclei matrix, and reduced dimensions with SVD (singular values = 50) similarly as for maize nuclei. We coded library sequence depth per nucleus as a covariate using the *dplyr* function *ntile* with n=3 and removed additional technical variance with *Harmony* using the SVD matrix as input with non-default settings for a weak correction (tau=3, nclust=15, max.iter.harmony=30, theta=0, lambda=10) (Korsunsky et al., 2019). Nuclei were clustered with Louvain clustering (resolution = 1) in the *Harmony* corrected embedding, and project into an additionally reduced space with UMAP (n_neighbors=15, min_dist=0.1).

#### Aligning pseudotime trajectories between A. thaliana and Z. mays

To enable comparison of companion cell development between *A. thaliana* and *Z. mays*, we first identified putative one-to-one orthologs using *OrthoFinder* (v2) (Emms and Kelly, 2019). Gene accessibility scores for 10,976 putative orthologs were imputed using a diffusion-based approach (Fang et al., 2020; van Dijk et al., 2018) and scaled from 0 to 1 across pseudotime for barcodes associated with companion cell development in *A. thaliana* and *Z. mays*. To account for different distributions, pseudotime coverage, and number of barcodes between species, we used the R package, *cellAlign,* that interpolates, scales, and weights gene accessibility scores on a fixed set of (n=200) equally spaced points (width parameter: winSz=0.1) from two trajectories to remove technical biases inherent to each data set(Alpert et al., 2018). For each putative ortholog, we performed global alignment of gene accessibility scores across *A. thaliana* and *Z. mays* pseudotime using the dynamic time warping algorithm with default settings in *cellAlign.* We then extracted the pseudotime shifts, representing the extent of gene accessibility deviation at any given point along the trajectory, for each putative ortholog. We clustered genes into two groups based on pseudotime shifts across companion cell development using k-means clustering. To identify conserved gene accessibility patterns across pseudotime, we clustered the normalized distances between *A. thaliana* and *Z. mays* putative orthologs using a mixture model (G=2) with the R package, *mclust.* The mixture model identified a bimodal distribution of normalized distances with 0.15 as a natural cut-off for defining conserved accessibility patterns. Putative orthologs with normalized distances less than the cut-off were placed in a third group defined as conserved. The above analysis was repeated with TF motif deviations scores for 440 TF motifs, without the need for ortholog searching as the same TF position weight matrices were used for both species, affording identical TF motif labels.

### ADDITIONAL RESOURCES

Cell-type resolved data can be viewed through our public Plant Epigenome JBrowse Genome Browser (Hofmeister and Schmitz, 2018) (http://epigenome.genetics.uga.edu/PlantEpigenome/index.html) by selecting either the *Z. mays* or *A. thaliana* Genome Browser links, followed by the scATAC_celltypes tab in the tracks panel.

